# Large-scale association analyses identify host factors influencing human gut microbiome composition

**DOI:** 10.1101/2020.06.26.173724

**Authors:** Alexander Kurilshikov, Carolina Medina-Gomez, Rodrigo Bacigalupe, Djawad Radjabzadeh, Jun Wang, Ayse Demirkan, Caroline I. Le Roy, Juan Antonio Raygoza Garay, Casey T. Finnicum, Xingrong Liu, Daria V. Zhernakova, Marc Jan Bonder, Tue H. Hansen, Fabian Frost, Malte C. Rühlemann, Williams Turpin, Jee-Young Moon, Han-Na Kim, Kreete Lüll, Elad Barkan, Shiraz A. Shah, Myriam Fornage, Joanna Szopinska-Tokov, Zachary D. Wallen, Dmitrii Borisevich, Lars Agreus, Anna Andreasson, Corinna Bang, Larbi Bedrani, Jordana T. Bell, Hans Bisgaard, Michael Boehnke, Dorret I. Boomsma, Robert D. Burk, Annique Claringbould, Kenneth Croitoru, Gareth E. Davies, Cornelia M. van Duijn, Liesbeth Duijts, Gwen Falony, Jingyuan Fu, Adriaan van der Graaf, Torben Hansen, Georg Homuth, David A. Hughes, Richard G. Ijzerman, Matthew A. Jackson, Vincent W.V. Jaddoe, Marie Joossens, Torben Jørgensen, Daniel Keszthelyi, Rob Knight, Markku Laakso, Matthias Laudes, Lenore J. Launer, Wolfgang Lieb, Aldons J. Lusis, Ad A.M. Masclee, Henriette A. Moll, Zlatan Mujagic, Qi Qibin, Daphna Rothschild, Hocheol Shin, Søren J. Sørensen, Claire J. Steves, Jonathan Thorsen, Nicholas J. Timpson, Raul Y. Tito, Sara Vieira-Silva, Uwe Völker, Henry Völzke, Urmo Võsa, Kaitlin H. Wade, Susanna Walter, Kyoko Watanabe, Stefan Weiss, Frank U. Weiss, Omer Weissbrod, Harm-Jan Westra, Gonneke Willemsen, Haydeh Payami, Daisy M.A.E. Jonkers, Alejandro Arias Vasquez, Eco J.C. de Geus, Katie A. Meyer, Jakob Stokholm, Eran Segal, Elin Org, Cisca Wijmenga, Hyung-Lae Kim, Robert C. Kaplan, Tim D. Spector, Andre G. Uitterlinden, Fernando Rivadeneira, Andre Franke, Markus M. Lerch, Lude Franke, Serena Sanna, Mauro D’Amato, Oluf Pedersen, Andrew D. Paterson, Robert Kraaij, Jeroen Raes, Alexandra Zhernakova

**Affiliations:** Department of Genetics, University of Groningen, University Medical Center Groningen, Groningen, the Netherlands; Department of Internal Medicine, Erasmus MC University Medical Center, Rotterdam, the Netherlands; The Generation R Study, Erasmus MC University Medical Center, Rotterdam, the Netherlands; Department of Microbiology and Immunology, Rega Instituut, KU Leuven, Leuven, Belgium; Center for Microbiology, VIB, Leuven, Belgium; Institute of Microbiology, Chinese Academy of Sciences, Beijing, China; Section of Statistical Multi-Omics, Department of Clinical & Experimental Medicine, School of Biosciences & Medicine, University of Surrey, Guildford, UK; Department of Twin Research & Genetic Epidemiology, King’s College London, London, UK; Department of Medicine, University of Toronto, Toronto, Canada; Division of Gastroenterology, Mount Sinai Hospital, Toronto, Canada; Avera Institute of Human Genetics, Avera McKennan Hospital & University Health Center, Sioux Falls, USA; Center for Molecular Medicine and Clinical Epidemiology Division, Department of Medicine Solna, Karolinska Institutet, Stockholm, Sweden; Laboratory of Genomic Diversity, Center for Computer Technologies, ITMO University, St. Petersburg, Russia; Novo Nordisk Foundation Center for Basic Metabolic Research, Faculty of Health and Medical Sciences, University of Copenhagen, Copenhagen, Denmark; Department of Medicine A, University Medicine Greifswald, Greifswald, Germany; Institute of Clinical Molecular Biology, Christian-Albrechts-University of Kiel, Kiel, Germany; Department of Epidemiology and Population Health, Albert Einstein College of Medicine, Bronx, USA; Medical Research Institute, Kangbuk Samsung Hospital, Sungkyunkwan University School of Medicine, Seoul, Republic of Korea; Department of Clinical Research Design and Evaluation, SAIHST, Sungkyunkwan University, Seoul, Republic of Korea; Estonian Genome Centre, Institute of Genomics, University of Tartu, Tartu, Estonia; Department of Computer Science and Cell Biology, Weizmann Institute of Science, Rehovot, Israel; COPSAC, Copenhagen University Hospital, Herlev-Gentofte, Copenhagen, Denmark; Institute of Molecular Medicine McGovern Medical School, The University of Texas Health Science Center at Houston, Houston, USA; Human Genetics Center School of Public Health, The University of Texas Health Science Center at Houston, Houston, USA; Department of Psychiatry, Radboudumc, Donders Institute for Brain, Cognition and Behaviour, Nijmegen, the Netherlands; Department of Neurology, University of Alabama at Birmingham, Birmingham, USA; Division of Family Medicine and Primary Care, Department of Neurobiology, Care Sciences and Society, Karolinska Institutet, Stockholm, Sweden; Stress Research Institute, Stockholm University, Stockholm, Sweden; Department of Biostatistics and Center for Statistical Genetics, University of Michigan, Ann Arbor, USA; Biological Psychology, Vrije Universiteit, Amsterdam, the Netherlands; Department of Pediatrics, Albert Einstein College of Medicine, Bronx, USA; Department of Microbiology & Immunology, Albert Einstein College of Medicine, Bronx, USA; Department of Epidemiology, Erasmus MC University Medical Center, Rotterdam, the Netherlands; Nuffield Department of Population Health, University of Oxford, Oxford, UK; Department of Pediatrics, Erasmus MC University Medical Center, Rotterdam, the Netherlands; Department of Pediatrics, University of Groningen, University Medical Center Groningen, Groningen, the Netherlands; Department of Functional Genomics, Interfaculty Institute for Genetics and Functional Genomics, University Medicine Greifswald, Greifswald, Germany; MRC Integrative Epidemiology Unit, University of Bristol, Bristol, UK; Population Health Sciences, Bristol Medical School, Bristol, UK; Department of Endocrinology, Amsterdam University Medical Center, location VUMC, Amsterdam, the Netherlands; Kennedy Institute of Rheumatology, University of Oxford, Oxford, UK; Centre for Clinical Research and Prevention, Bispebjerg/Frederiksberg Hospital, Capital Region of Copenhagen and Faculty of Health and Medical Sciences, University of Copenhagen, Copenhagen, Denmark; Division of Gastroenterology-Hepatology, Maastricht University Medical Center+, Maastricht, the Netherlands; NUTRIM School of Nutrition and Translational Research in Metabolism, Maastricht University, Maastricht, the Netherlands; Department of Pediatrics, University of California San Diego, La Jolla, USA; Center for Microbiome Innovation, University of California San Diego, La Jolla, USA; Center for Microbiome Innovation and department of Bioengeering, University of California San Diego, La Jolla, USA; Institute of Clinical Medicine, Internal Medicine, University of Eastern Finland, Kuopio, Finland; Department of Medicine I, University Hospital Schleswig-Holstein, Campus Kiel, Kiel, Germany; Laboratory of Epidemiology and Population Science, National Institute on Aging, Bethesda, USA; Institute of Epidemiology, Kiel University, Kiel, Germany; Departments of Microbiology, Immunology and Molecular Genetics, and Human Genetics, University of California, Los Angeles, Los Angeles, USA; Department of Medicine, University of California, Los Angeles, Los Angles, USA; Department of Family Medicine, Kangbuk Samsung Hospital, Sungkyunkwan University School of Medicine, Seoul, Republic of Korea; Center for Cohort Studies, Total Healthcare Center, Kangbuk Samsung Hospital, Sungkyunkwan University School of Medicine, Seoul, Republic of Korea; Department of Biology, University of Copenhagen, Copenhagen, Denmark; Institute for Community Medicine, University Medicine Greifswald, Greifswald, Germany; Department of Biomedical and Clinical Sciences, University of Linköping, Linköping, Sweden; Department of gastroenterology, County Council of Östergötland, Linköping, Sweden; Department of Complex Trait Genetics, Center for Neurogenomics and Cognitive Research, Neuroscience Campus Amsterdam, VU University Amsterdam, Amsterdam, the Netherlands; School of Public Health, Harvard University, Boston, USA; Department of Human Genetics, Radboudumc, Donders Institute for Brain, Cognition and Behaviour, Nijmegen, the Netherlands; Amsterdam Public Health, Amsterdam UMC, Amsterdam, the Netherlands; Department of Nutrition, University of North Carolina at Chapel Hill, Chapel Hill, USA; Nutrition Research Institute, University of North Carolina at Chapel Hill, Kannapolis, USA; Department of Biochemistry, Ewha Womans University School of Medicine, Seoul, Republic of Korea; Division of Public Health Sciences, Fred Hutchinson Cancer Research Center, Seattle, USA; Istituto di Ricerca Genetica e Biomedica, National Research Council, Monserrato, Italy; School of Biological Sciences, Monash University, Clayton, Australia; Department of Gastrointestinal and Liver Diseases, Biodonostia Health Research Institute, San Sebastián, Spain; Ikerbasque, Basque Science Foundation, Bilbao, Spain; Genetics and Genome Biology, The Hospital for Sick Children Research Institute, Toronto, Canada

## Abstract

To study the effect of host genetics on gut microbiome composition, the MiBioGen consortium curated and analyzed genome-wide genotypes and 16S fecal microbiome data from 18,340 individuals (24 cohorts). Microbial composition showed high variability across cohorts: only 9 out of 410 genera were detected in more than 95% samples. A genome-wide association study (GWAS) of host genetic variation in relation to microbial taxa identified 31 loci affecting microbiome at a genome-wide significant (P<5×10^−8^) threshold. One locus, the lactase (*LCT*) gene locus, reached study-wide significance (GWAS signal P=1.28×10^−20^), and it showed an age-dependent association with *Bifidobacterium* abundance. Other associations were suggestive (1.95×10^−10^<P<5×10^−8^) but enriched for taxa showing high heritability and for genes expressed in the intestine and brain. A phenome-wide association study and Mendelian randomization identified enrichment of microbiome trait loci in the metabolic, nutrition and environment domains and suggested the microbiome has causal effects in ulcerative colitis and rheumatoid arthritis.

## Introduction

The gut microbiome is an integral part of the human holobiont. In recent years, many studies have highlighted the link between its perturbations and immune, metabolic, neurologic and psychiatric traits, drug metabolism and cancer^1^. Environmental factors, like diet and medication, play a significant role in shaping the gut microbiome composition^2–4^, although twin, family and population-based studies have shown that the genetic component also plays a role in determining gut microbiota composition, and a proportion of bacterial taxa are heritable^5,6^.

Several studies^7–9^ have investigated the effect of genetics on microbiome composition through genome-wide association studies (GWAS) and identified dozens of associated loci. However, little cross-replication across these studies has been observed so far^10,11^. This may be due to a number of factors. First, methodological differences in the collection, processing and annotation of stool microbiota are known to have significant effects on the microbiome profiles obtained^12–14^ and can generate heterogeneity and a lack of reproducibility across studies. Second, most association signals are rather weak, which suggests that existing studies of 1,000–2,000 samples^7–9^ are underpowered. Finally, some of the GWAS signals related to microbiome compositions may be population-specific, i.e. they may represent *bona fide* population differences in genetic structure and/or environment.

To address these challenges and obtain valuable insights into the relationship between host genetics and microbiota composition, we set up the international consortium MiBioGen^11^. In this study, we have coordinated 16S rRNA gene sequencing profiles and genotyping data from 18,340 participants from 24 cohorts from the USA, Canada, Israel, South Korea, Germany, Denmark, the Netherlands, Belgium, Sweden, Finland and the UK. We performed a large-scale, multi-ethnic, genome-wide meta-analysis of the associations between autosomal human genetic variants and the gut microbiome. We explored the variation of microbiome composition across different populations and investigated the effects of differences in methodology on the microbiome data. Through the implementation of a standardized pipeline, we then performed microbiome trait loci (mbTL) mapping to identify genetic loci that affect the relative abundance (mbQTLs) or presence (microbiome Binary Trait loci, or mbBTLs) of microbial taxa. Finally, we focused on the biological interpretation of GWAS findings through Gene Set Enrichment Analysis (GSEA), Phenome-wide association studies (PheWAS) and Mendelian randomization (MR) approaches.

## Results

### Landscape of microbiome composition across cohorts

Our study included cohorts that were heterogeneous in terms of ethnic background, age, male/female ratio and microbiome analysis methodology. Twenty cohorts included samples of single ancestry, namely European (16 cohorts, N=13,266), Middle-Eastern (1 cohort, N=481), East Asian (1 cohort, N=811), American Hispanic/Latin (1 cohort N=1,097) and African American (1 cohort, N=114), whereas four cohorts were multi-ancestry (N=2,571) (see Supplementary Note, Supplementary Tables 1,2).

Twenty-two cohorts comprised adult or adolescent individuals (N=16,632), and two cohorts consisted of children (N=1,708). The microbial composition was profiled by targeting three distinct variable regions of the 16S rRNA gene: V4 (10,413 samples, 13 cohorts), V3-V4 (4,211 samples, 6 cohorts) and V1-V2 (3,716 samples, 5 cohorts) (Fig. 1a). To account for differences in sequencing depth, all datasets were rarefied to 10,000 reads per sample. Next, we performed taxonomic classification using direct taxonomic binning instead of OTU clustering methods (see Online Methods)^11,15,16^.

**Figure 1.**
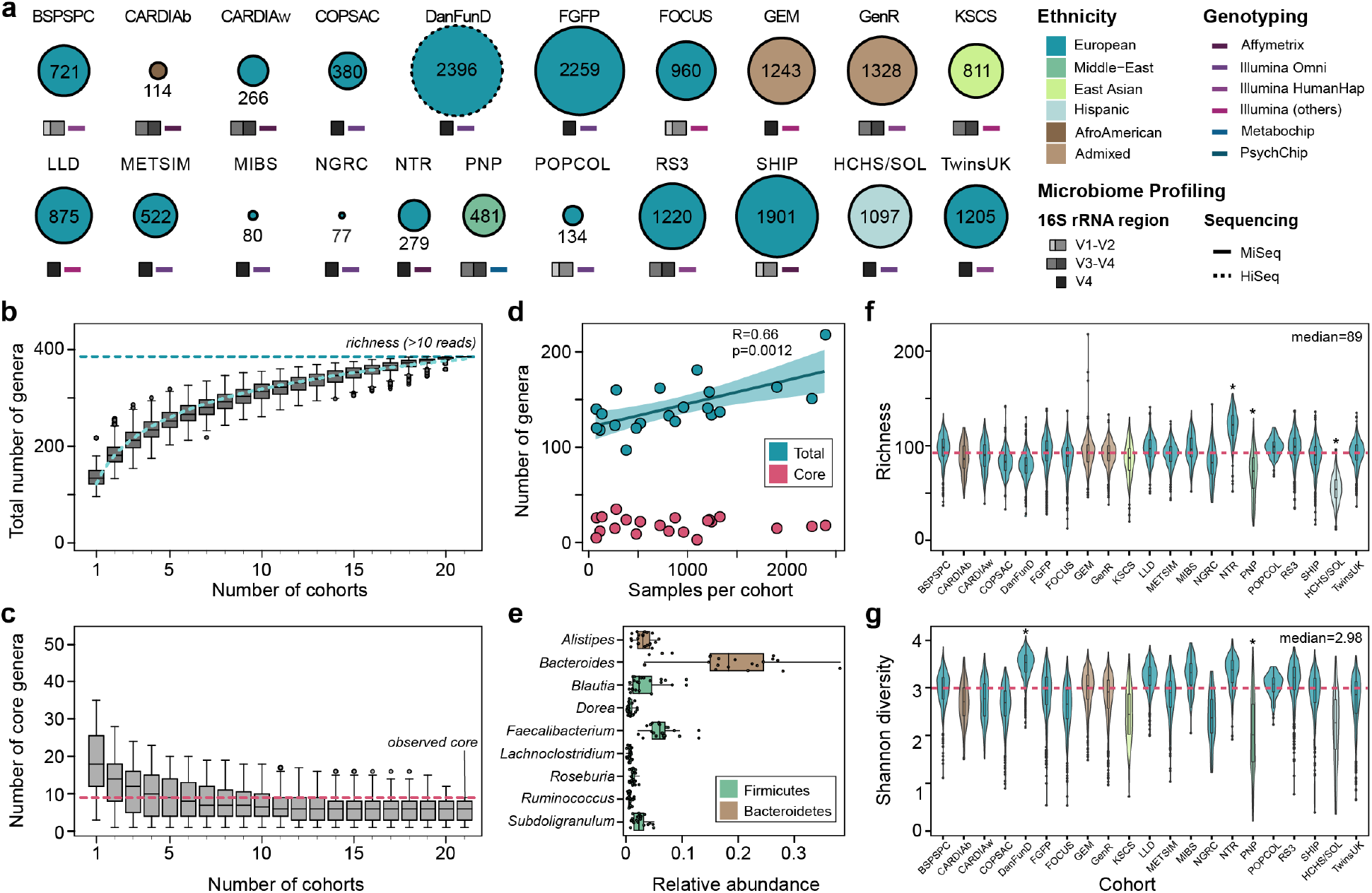
Diversity of microbiome composition across the MiBioGen cohorts. **(a)** Sample size, ethnicity, genotyping array and 16S rRNA gene profiling method. The SHIP/SHIP-TREND and GEM_v12/GEM_v24/GEM_ICHIP subcohorts are combined in SHIP and GEM, respectively (Online Methods; see Supplementary Note for cohort abbreviations). This merge resulted in the total of 21 cohorts depicted in the figure. **(b)\*** Total richness (number of genera with mean abundance over 0.1%, i.e. 10 reads out of 10,000 rarefied reads) by number of cohorts investigated. **(c)\*** Number of core genera (genera present in >95% of samples from each cohort) by number of cohorts investigated. **(d)** Pearson correlation of cohort sample size with total number of genera. Confidence band represents the standard error of the regression line. **(e)\*** Unweighted mean relative abundance of core genera across the entire MiBioGen dataset. **(f)\*** Per-sample richness across the 21 cohorts. Asterisks indicate cohorts that differ significantly from all the others (pairwise Wilcoxon rank-sum test; FDR<0.05). **(g)** Diversity (Shannon index) across the 21 cohorts, with the DanFund and PNP cohorts presenting higher and lower diversity in relation to the other cohorts (pairwise Wilcoxon rank sum test; FDR<0.05). **(*)** For all boxplots, the central line, box and whiskers represent the median, IQR and 1.5 times the IQR.

In general, cohorts varied in their microbiome structure at multiple taxonomic levels (Fig. 1b-g). This variation may largely be driven by the heterogeneity between populations and differences in technical protocols (Supplementary Tables 1-3). Combining all samples (N=18,340) resulted in a total richness of 385 genus-level taxonomic groups that had a relative abundance higher than 0.1% in at least one cohort. This observed total richness appears to be below the estimated saturation level (Fig. 1b), suggesting that a further increase in sample size and a higher sequencing depth are needed to capture the total gut microbial diversity (Fig. 1d). As expected, the core microbiota (the number of bacterial taxa present in over 95% of individuals) decreased with the inclusion of additional cohorts (Fig. 1c, Online Methods). The core microbiota comprise nine genera, of which seven were previously identified as such^3^, plus the genera *Ruminococcus* and *Lachnoclostridium* (Fig. 1e). Of these nine genera, the most abundant genus was *Bacteroides* (18.65% (SD:8.65)), followed by *Faecalibacterium* (6.19% (SD:2.35)), *Blautia* (3.36% (SD:2.84)) and *Alistipes* (3.05% (SD:1.47)). Among the European cohorts that compose the largest genetically and environmentally homogeneous cluster, the core microbiota also included *Ruminiclostridium, Fusicatenibacter, Butyricicoccus* and *Eubacterium*, genera which typically produce short-chain fatty acids^17^.

The DNA extraction method was the principal contributor to heterogeneity, with a non-redundant effect size of 29% on the microbiome variation (measured as average genus abundance per cohort; stepwise distance-based redundancy analysis R2adj_DNAext_=0.27, P_adj_=7×10^−4^) (Supplementary Table 4). Richness and Shannon diversity also differed significantly across cohorts. The cohorts with the lowest richness (HCHS/SOL) and highest diversity (DanFund) used specific DNA extraction kits that were not used by other studies, possibly contributing to their outlying alpha diversities (Fig. 1f,g, Supplementary Table 3). Overall, the 16S rRNA domain sequenced and the DNA extraction methods used, together with cohort ethnicity, accounted for 32.74% of richness variance.

Given the high heterogeneity of microbial composition across cohorts, we applied both per-cohort and whole study–filters for taxa inclusion in GWAS (see Online Methods).

### Heritability of microbial taxa and alpha diversity

We performed estimation of heritability (H^2^) of gut microbiome composition based on the two twin cohorts included in our study (Supplementary Table 5). The TwinsUK cohort, composed of 1,176 samples, including 169 monozygotic (MZ) and 419 dizygotic (DZ) twin pairs, was used to estimate H^2^ using the ACE (additive genetic variance (A)/shared environmental factors (C)/ non-shared factors plus error (E)) model. The Netherlands Twin Registry (NTR) cohort (only MZ twins, N=312, 156 pairs) was used to replicate the MZ intraclass correlation coefficient (ICC). None of the alpha diversity metrics (Shannon, Simpson and inverse Simpson) showed evidence for heritability (A<0.01, P=1). Among the 159 bacterial taxa that were present in more than 10% of twin pairs, 19 taxa showed evidence for heritability (P_nominal_<0.05) (Fig. 2a). The ICC shows concordance between TwinsUK and NTR for these 19 bacterial taxa (R=0.25, P=0.0018, Fig. 2b).

**Figure 2.**
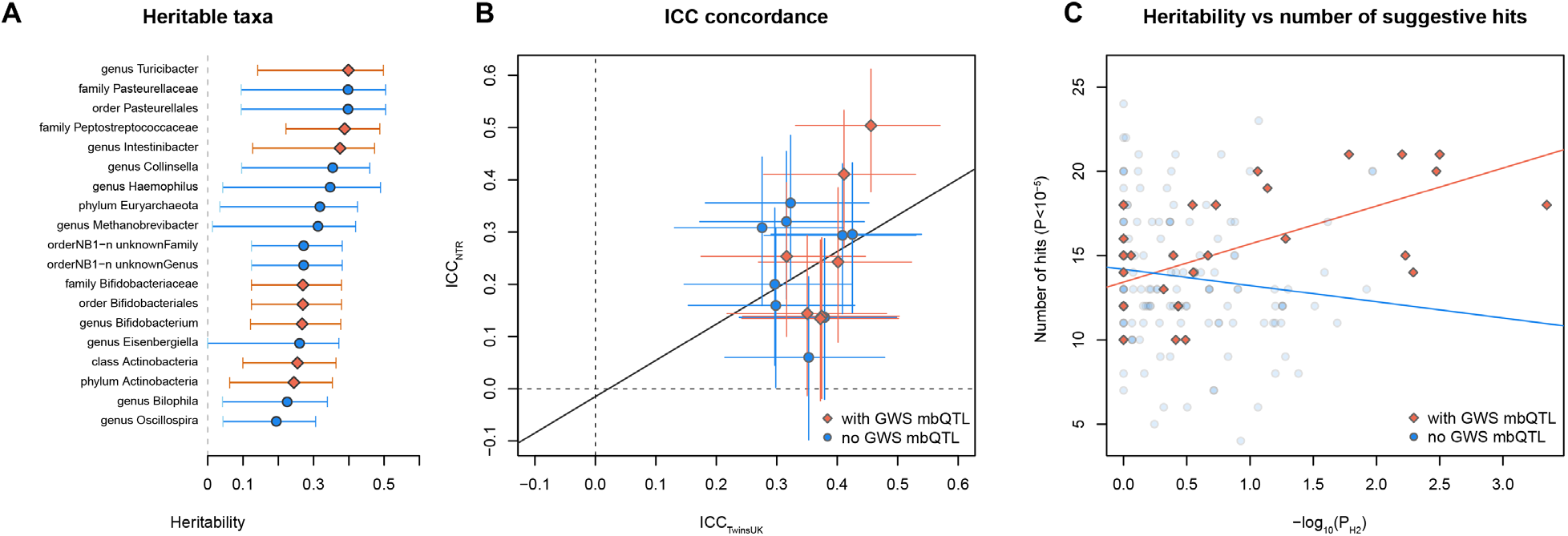
Heritability of microbiome taxa and its concordance with mbQTL mapping. **(a)** Microbial taxa that showed significant heritability in the TwinsUK cohort (ACE model, nominal P<0.05, no adjustment for multiple comparison). Taxa with at least one genome-wide significant (GWS) mbQTL hit are marked red. Only taxa present in more than 10% of pairs (>17 MZ pairs, >41 DZ pairs) are shown. Circles and diamonds represent heritability value. Error bars represent 95% CI. **(b)** Correlation of monozygotic ICC between TwinsUK and NTR cohort. Only taxa with significant heritability (ACE model P<0.05) that are present in both TwinsUK and NTR are shown. Red and blue dots indicate bacterial taxa with/without GWS mbQTLs (P<5×10^−8^), respectively. Segments represent 95% CI. **(c)** Correlation between heritability significance (-log_10_P_H2_ TwinsUK) and the number of loci associated with microbial taxon at relaxed threshold (P_mbQTL_<1×10^−5^). Taxa with at least one GWS-associated locus are marked red. Error bars represent 95% confidence intervals.

The SNP-based heritability calculated from mbQTL summary statistics using linkage disequilibrium (LD) score regression showed two bacterial taxa, genus *Ruminiclostridium* 9 and family *<u>Peptostreptococcaeae</u>*, passing the significance threshold given the number of 211 taxa tested (Z<3.68, Supplementary Table 5). The results of the SNP-based heritability and twin-based heritability showed significant correlation across the tested taxa (R=0.244, P=7.2×10^−4^).

### Thirty one loci associated with gut microbes through GWAS

First, we studied the genetic background of the alpha diversity (Simpson, inverse Simpson and Shannon diversity indices). We identified no significant hits in the meta-GWAS (P>5×10^−8^; Supplementary Table 6, Supplementary Fig. 1), which is in line with the observed lack of heritability for these indices.

Next, we used two separate GWAS meta-analysis approaches^18–20^ to explore the effect of host genetics on the abundance levels (mbQTL) or presence/absence (mbBTL) of bacterial taxa in the gut microbiota (see Online Methods).

In total, 18,340 samples and 211 taxa were included in the mbQTL mapping analysis (Online Methods, Supplementary Table 3). We identified genetic variants that mapped to 20 distinct genetic loci associated with the abundance of 27 taxa (Fig. 3; Supplementary Fig. 2,3; Supplementary Table 7, 8). MbBTL mapping covered 177 taxa, and 10 loci were found to be associated with presence/absence of bacterial taxa (Fig. 3, Supplementary Table 7, 9). For one taxon, family *Peptococcaceae*, two independent mbBTLs were detected (Fig. 3, Supplementary Table 7). Two out of 30 mbTLs showed heterogeneity in mbTL effect-sizes (Supplementary Note).

**Figure 3.**
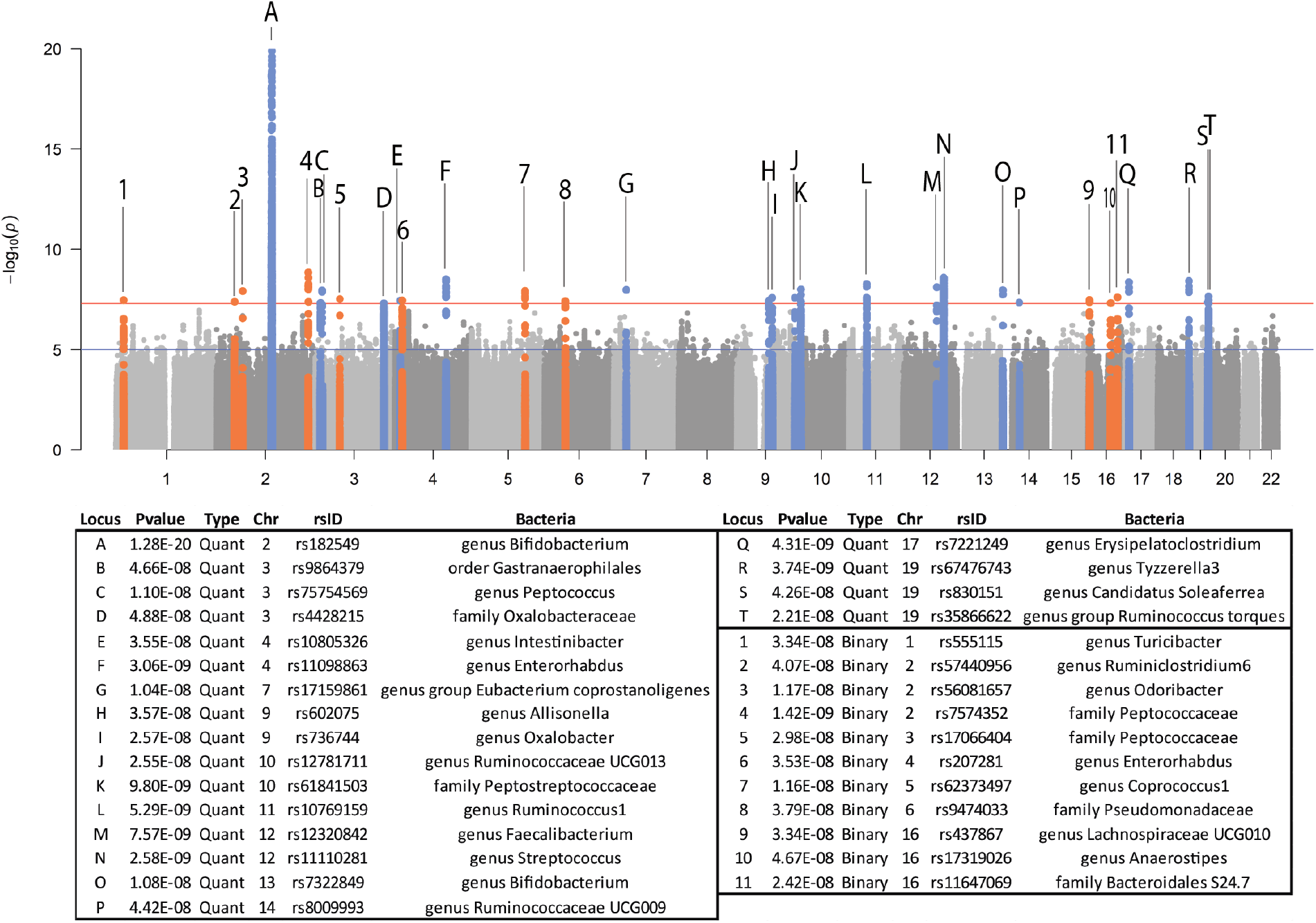
Manhattan plot of the mbTL mapping meta-analysis results. MbQTLs are indicated by letters. MbBTLs are indicated by numbers. For mbQTLs, the Spearman correlation test (two-sided) was used to identify loci that affect the covariate-adjusted abundance of bacterial taxa, excluding samples with zero abundance. For mbQTLs, p-values (two-sided) were calculated by logistic regression. Horizontal lines define nominal genome-wide significance (P=5×10^−8^, red) and suggestive genome-wide (P=1×10_-5_, blue) thresholds.

In both the mbQTL and mbBTL mapping, only one out of 31 loci (*LCT* locus – *Bifidobacterium*, P=8.63×10^−21^) passed the strict correction for the number of taxa tested (P<1.95×10^−10^ for 257 taxa included in the analysis). However, the remaining loci include functionally relevant variants (i.e. the *FUT2* gene suggested by earlier studies^21^) and, overall, showed concordance with the heritability of microbial taxa. Seven out of the nine taxa that showed the strongest evidence for heritability in the TwinsUK cohort (P<0.01) also have genome-wide significant mbTLs (Fig. 2b). For the taxa with genome-wide significant mbTLs, the number of independent loci associated with a relaxed threshold of 1×10^−5^ strongly correlated with heritability significance (R=0.62, P=1.9×10^−4^, Fig. 2c), suggesting that more mbTLs would be identified for this group of bacteria using a larger sample size.

### *LCT* mbQTL effect shows age and ethnic heterogeneity

The strongest association signal was seen for variants located in a large block of about 1.5Mb at 2q21.3, which includes the *LCT* gene and 12 other protein-coding genes. This locus has previously been associated with the abundance of *Bifidobacterium* in Dutch^7^, UK^6^ and US^22^ cohorts. Previous studies have also shown a positive correlation of *Bifidobacterium* abundance with the intake of milk products, but only in individuals homozygous for the low-function LCT haplotype, thereby indicating that gene–diet interaction regulates *Bifidobacterium* abundance^7^. In our study, the strongest association was seen for rs182549 (P=1.28×10^−20^), which is a perfect proxy for the functional *LCT* variant rs4988235 (r^2^=0.996, D’=1 in European populations). This association showed evidence for heterogeneity across cohorts (I^2^=62.73%, Cochran’s Q P=1.4×10^−4^). A leave-one-out strategy showed that the COPSAC_2010_ cohort, which includes children 4-6 years of age range, contributed the most to the detected heterogeneity (Fig. 4a,b; Supplementary Table 2). When this study was excluded from the meta-analysis, the heterogeneity was reduced (I^2^=51.9%, Cochran’s Q P=0.004). A meta-regression analysis showed that linear effects of age and ethnicity accounted for 11.84% of this heterogeneity. Including quadratic and cubic terms of age in the model explained 39.22% of the heterogeneity, and the residual heterogeneity was low (Cochran’s Q P=0.01) (Fig. 4c).

**Figure 4.**
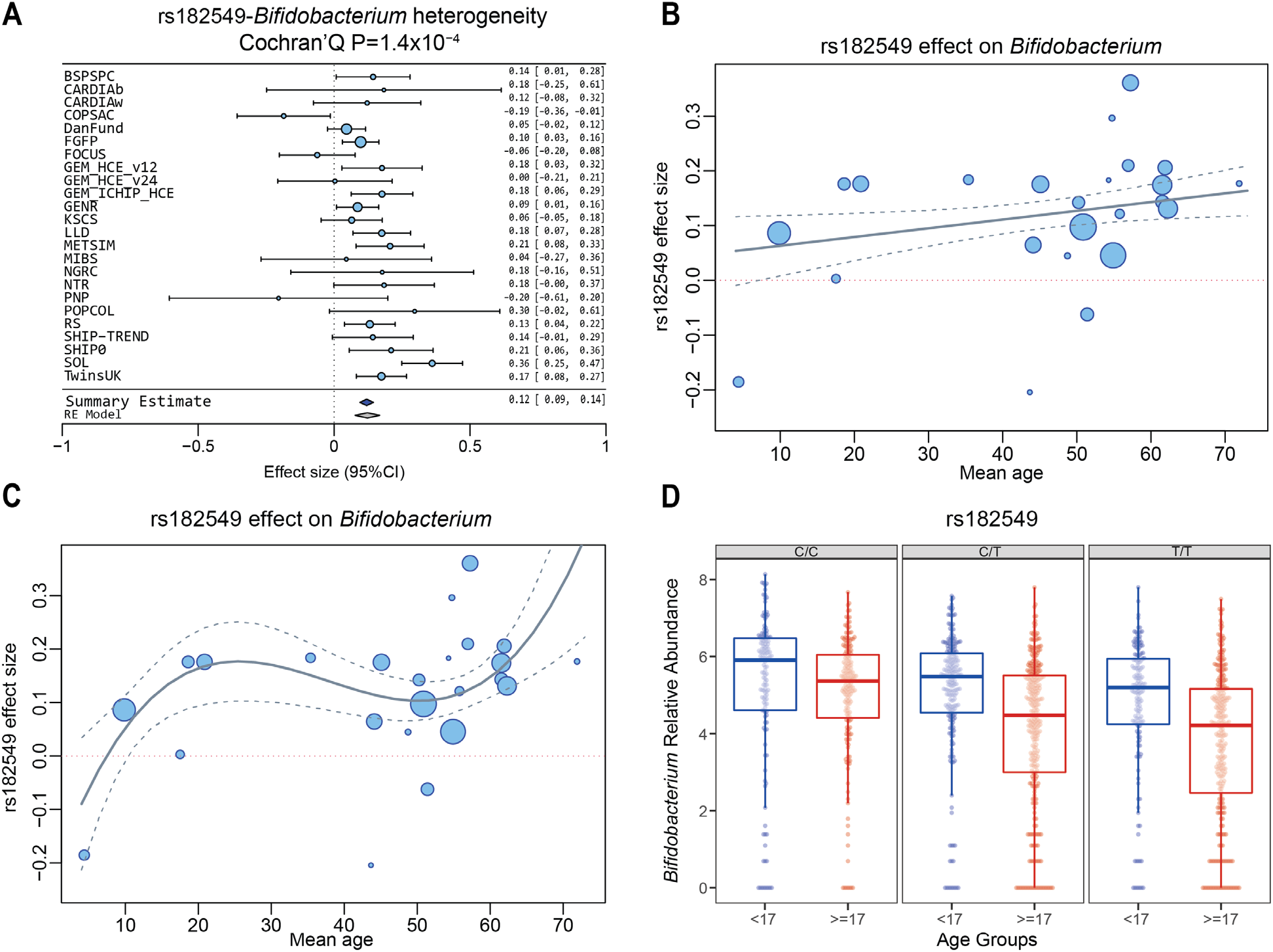
Association of the *LCT* locus (rs182549) with the genus *Bifidobacterium*. **(a)** Forest plot of effect sizes of rs182549 and abundance of *Bifidobacterium*. Effect sizes and 95% CI are defined as circles and error bars. Effect sizes were calculated from Spearman correlation p-values (Online Methods). **(b)** Meta-regression of the association of mean cohort age and mbQTL effect size. Confidence bands represent the standard error of the meta-regression line. **(c)** Meta-regression analysis of the effect of linear, squared and cubic terms of age on mbQTL effect size. Confidence bands represent the standard error of the meta-regression line. **(d)** Age-dependence of mbQTL effect size in the GEM cohort. Blue boxes include samples in the age range 6–16 years old. Red boxes include samples with age ≥17 years. The C/C (rs182549) genotype is a proxy of the NC_000002.11:g.136608646= (rs4988235) allele, which is associated to functional recessive hypolactasia. The central line, box and whiskers represent the median, IQR and 1.5 times the IQR, respectively. See Supplementary Note for cohort abbreviations.

Following these observations, we decided to investigate the effect of age and ethnicity in the multi-ethnic GEM cohort, comprising 1,243 individuals with an age range between 6 and 35 years, of which nearly half of the participants are 16 years or younger. Our analysis showed a significant SNP–age interaction on the level of *Bifidobacterium* abundance (P<0.05, see Online Methods). Individuals homozygous for the NC_000002.11:g.136616754CC (rs182549) genotype showed a higher abundance of the genus *Bifidobacterium* in the adult group, but not in the younger group (Fig. 4d). The age–genotype interaction was significant in the GEM_v12 and GEM_ICHIP subcohorts, both comprising mostly European individuals, while the GEM_v24 cohort, mainly composed of individuals of different Israeli subethnicities (see Online Methods) who live in Israel, showed neither an mbQTL effect (Beta = -0.002 [95%CI: -0.21, 0.21]) nor an interaction with age (P>0.1). The lack of an *LCT* mbQTL effect in adults was also observed in another Israel cohort in the study (Personalized Nutrition Project (PNP), 481 adults, Beta = -0.20 [95%CI: -0.61, 0.20]). Altogether, the cohorts that reported the lowest *LCT* effect sizes were the two cohorts of Israeli ethnicity volunteered in Israel (GEM_v24, PNP) and a child cohort (Copenhagen Prospective Studies on Asthma in Childhood (COPSAC), Beta = -0.18 [95%CI: - 0.36, -0.01]).

### mbTLs are enriched for genes related to metabolism

Several loci detected at genome-wide significance level were enriched for genes related to metabolism.

In the mbQTL analysis, the *FUT2-FUT1* locus was associated to the abundance of the *Ruminococcus torques* genus group, a genus from the *Lachnospiraceae* family. The leading SNP (rs35866622 for *R. torques* group, P=2.21×10^−8^) is a proxy for the functional variant rs601338 (r^2^=0.8; D’=0.9 in European populations) that introduces a stop-codon in *FUT2*^23^. Another proxy of the functional *FUT2* SNP, rs281377, showed association to the *Ruminococcus gnavus* genus group in the binary analysis, however this signal was just above the genome-wide significance threshold (P=5.79×10^−8^) (Supplementary Table 9). *FUT2* encodes the enzyme alpha-1,2-fucosyltransferase, which is responsible for the secretion of fucosylated mucus glycans in the gastrointestinal mucosa^24^. Individuals homozygous for the stop-codon (rs601338*A/A, non-secretors) do not express ABO antigens on the intestinal mucosa. We observed that the tagging NC_000019.9:g.49218060C>T (rs35866622 non-secretor) allele was associated with a reduced abundance of the *R. torques* group and a decreased presence of the *R. gnavus* group. *Ruminococcus sp*. are specialized in the degradation of complex carbohydrates^25^, thereby supporting a link between genetic variation in the *FUT2* gene, levels of mucus glycans and the abundance of this taxa. When assessing the link between this variant and phenotypes in the LifeLines-DEEP (LLD; N=875) and Flemish Gut Flora Project (FGFP, N=2,259) cohorts (Online Methods), the strongest correlation for the *R. torques* group was seen with fruit intake (LLD: R_Sp_=-0.19, P_adj_=3.1×10^−5^; FGFP: R_Sp_=-0.10, P_adj_=1.4×10^−4^, Supplementary Table 10, 11), in line with the association of *FUT2* with food preferences, as discussed in the results of the PheWAS (see below).

Several other suggestive mbQTLs can be linked to genes potentially involved in host– microbiome crosstalk. One of them includes three SNPs in 9q21 (top-SNP rs602075, P=3.57×10^− 8^) associated with abundance of *Allisonella*. The 9q21 locus includes the genes *PCSK5, RFK* and *GCNT1*, of which *RFK* encodes the enzyme that catalyzes the phosphorylation of riboflavin (vitamin B2) and *GCNT1* encodes a glycosyltransferase involved in biosynthesis of mucin. These products play major roles in the host–microbiota interactions within the intestine, where they are used by bacteria for their metabolism and involved in the regulation of the host immune defense^26^. Another association signal 10p13 (rs61841503, P=9.8×10^−9^), which affects the abundance of the heritable family *Peptostreptococcaceae*, is located in the *CUBN* gene, the receptor for the complexes of cobalamin (vitamin B12) with gastric intrinsic factor (the complex required for absorption of cobalamin). *CUBN* is expressed in the kidneys and the intestinal epithelium and is associated with B12-deficient anemia and albuminuria^27^. Cobalamin is required for host–microbial interactions^28^, and supplementation with cobalamin induced a substantial shift in the microbiota composition of an *in vitro* colon model^29^. These associations suggest that some members of the gut microbiome community might be affected by genetic variants that regulate the absorption and metabolism of vitamins B2 and B12.

Among mbBTLs, the strongest evidence for association was seen for a block of 10 SNPs (rs7574352, P=1.42×10^−9^) associated with the family *Peptococcaceae*, a taxon negatively associated with stool levels of the gut inflammation markers chromogranin A (LLD: R_Sp_=-0.31, P_adj_=4.4×10^−18^, Table S10) and calprotectin (LLD: R_Sp_=-0.11, P_adj_=0.058) and with ulcerative colitis (FGFP: R_Sp_=-0.06, P_adj_=0.09, Table S11). The association block is located in the intergenic region in the proximity (220kb apart) of *IRF1*, which is involved in insulin resistance and susceptibility to type 2 diabetes^30^.

Other highlights of identified mbTLs are given in the Supplementary Note.

### GSEA, FUMA and PheWAS analysis

To explore the potential functions of the identified mbTLs, we performed FUMA (Functional Mapping and Annotation of GWAS, see Online Methods)^31^, GSEA and PheWAS, followed by Bayesian colocalization analysis and genetic correlation of *Bifidobacterium* abundance to its PheWAS-related traits. FUMA of 20 mbQTL loci returned 139 positional and eQTL genes. GSEA on these genes suggested an enrichment for genes expressed in the small intestine (terminal ileum) and brain (*substania nigra* and putamen basal ganglia) (Supplementary Fig. 4). The positional candidates for mbBTLs did not show any enrichment in GSEA analysis.

To systematically assess the biological outcomes of the mbTLs, we looked up the 31 mbTLs in the summary statistics for 4,155 complex traits and diseases using the GWAS ATLAS^32^. Five out of 31 leading SNPs were associated with one or more phenotypes at P<5×10^− 8^ (Supplementary Table 12): rs182549 (*LCT*) and rs35866622 (*FUT1/ FUT2*), followed by rs4428215 (*FNDC3B*), rs11647069 (*PM FBP1)* and rs9474033 (*PKHD1*).

The variant showing highest pleiotropy, rs182549 (*LCT, Bifidobacterium*), was associated with multiple dietary and metabolic phenotypes, and the causal involvement of the SNP across pairs of traits was confirmed by colocalization test (PP.H4.abf > 0.9) for 49 out of 51 tested phenotypes. The NC_000002.11:g.136616754= (rs182549) allele, which predisposes individuals to lactose intolerance, was negatively associated with obesity^33^ and positively associated with Type 2 diabetes mellitus (T2DM) diagnosis (OR=1.057 [95%CI:1.031, 1.085], P=1.74×10^−5^), family history of T2DM (paternal: OR=1.054 [95%CI:1.035, 1.073], P=1.41×10^−8^; maternal: OR=1.035 [95%CI:1.016, 1.053], P=0.0002, siblings: OR=1.03 [95%CI:1.009, 1.052]), and several nutritional phenotypes in the UK Biobank cohort^32^. Moreover, the functional *LCT* SNP rs4988235 variant is associated with 1,5-anhydroglucitol (P=4.23×10^−28^)^34^, an indicator of glycemic variability^35^. There was a nominally significant correlation of *Bifidobacterium* with raw vegetable intake (rg=0.36, P=0.0016), but this correlation was not statistically significant after correction for multiple testing.

NC_000019.9:g.49218060= (rs35866622, *FUT1*/*FUT2* locus) was positively associated with fish intake and height. The secretor allele was negatively associated with the risks of cholelithiasis and Crohn’s disease, alcohol intake frequency, high cholesterol and waist-to-hip ratio (adjusted for body mass index (BMI), with PP.H4.abf > 0.9).

Consistent with the single SNP analysis, gene-based PheWAS also showed a strong link of the *LCT* locus with metabolic traits (e.g. P=5.7×10^−9^ for BMI), whereas several nutritional (e.g. P=1.26×10^−20^ for oily fish intake), immune-related (e.g. P=1.73×10^−12^ for mean platelet volume), gastrointestinal (e.g. P=8.77×10^−14^ for cholelithiasis) and metabolic signals (e.g. P=1.13×10^−13^, high cholesterol) mapped to the *FUT1*/*FUT2* locus (Fig. 5, Supplementary Table 13).

**Figure 5.**
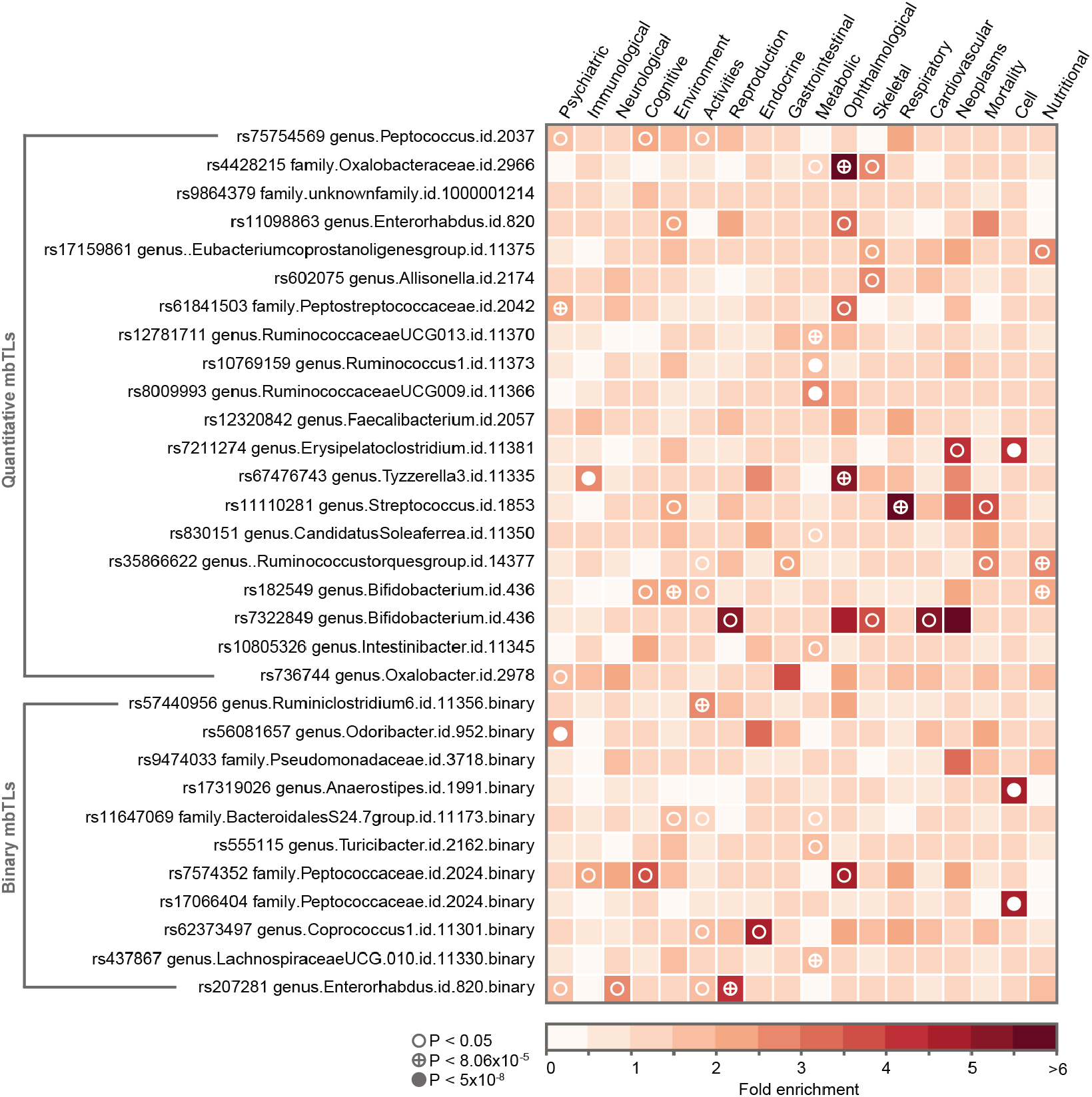
Phenome-wide association study (PheWAS) domain enrichment analysis. The analysis covered top-SNPs from 30 mbTLs and 20 phenotype domains. Three thresholds for multiple testing were used: 0.05, 8.3×10^−5^ (Bonferroni adjustment for number of phenotypes and genotypes studied) and 5×10^−8^ (an arbitrary genome-wide significance threshold). Only categories with at least one significant enrichment signal are shown.

Finally, we performed a phenotype domain enrichment analysis (Online Methods). We observed that top loci were enriched with signals associated with the metabolic domain supported by 4 mbTLs, followed by nutritional, cellular, immunological, psychiatric, ophthalmological, respiratory and reproductive traits, and the activities domain (Fig. 5, Supplementary Table 14).

### Mendelian Randomization analysis

To identify the potential causal links between gut microbial taxa and phenotypes, we performed bi-directional two-sample MR analyses using the TwoSampleMR package^36^. We focused on two groups of phenotypes: diseases (autoimmune, cardiovascular, metabolic and psychiatric) and nutritional phenotypes^37–42^. The complexity of the mechanisms by which host genetics affect microbiome composition, and the limited impact of genetic variants on microbial taxa variability, require caution when performing and interpreting causality estimation using MR analysis^43^. We therefore carried out several sensitivity analyses and excluded any results that showed evidence of being confounded by pleiotropy (Online Methods). Only pairs supported by three or more SNPs were considered. With these strict cut-offs, no evidence for causal relationships between microbiome taxa and dietary preferences was identified (Supplementary Tables 15, 16). However, our results suggest that a higher abundance of the class Actinobacteria and its genus *Bifidobacterium* may have a protective effect on ulcerative colitis (Actinobacteria: OR=0.56 [95%CI: 0.44-0.71] for each SD increase in bacterial abundance, P_BHadj_=8.8×10^−4^; *Bifidobacterium*: OR=0.51 [95%CI: 0.39-0.71], P_BHadj_=9.8×10^−5^) (Fig. 6a,b). We also observed that higher abundance of family *Oxalobacteraceae* has a protective effect on rheumatoid arthritis (OR=0.82, [95%CI: 0.74-0.91], P_BHadj_=0.028, Fig. 6c).

**Figure 6.**
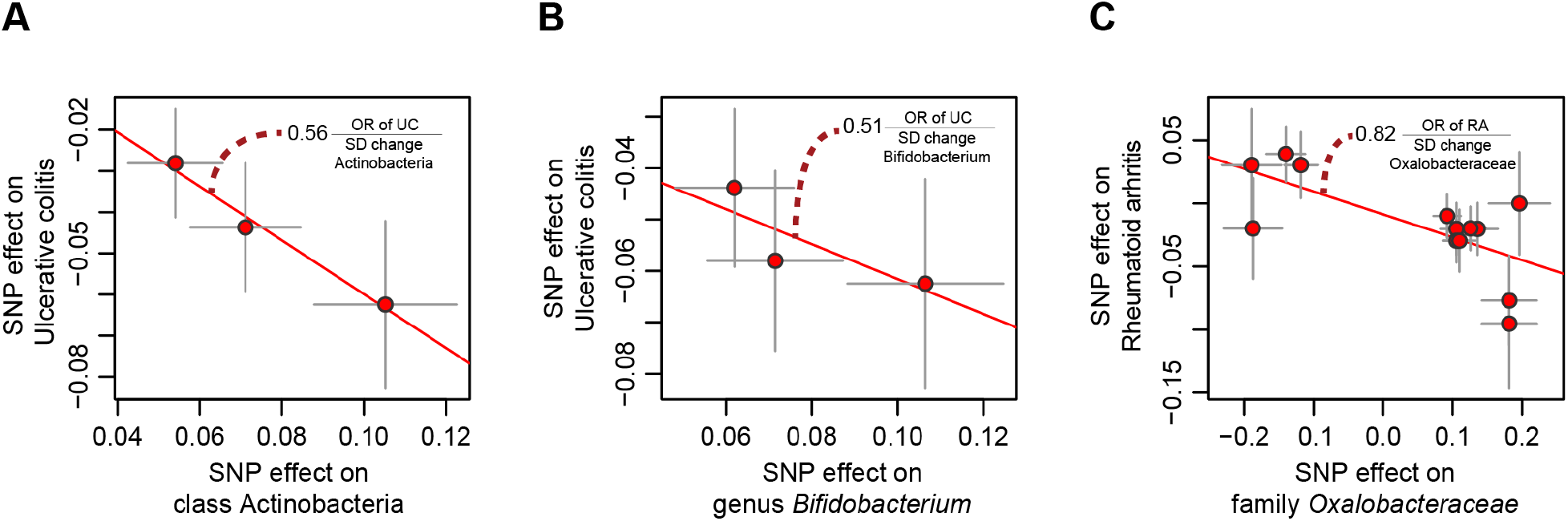
Mendelian randomization (MR) analysis. The X-axes show the SNP-exposure effect and the Y-axes show the SNP-outcome effect (SEs denoted as segments). **(a)** MR analysis of class Actinobacteria (exposure) and ulcerative colitis (outcome). **(b)** MR analysis of genus *Bifidobacterium* (exposure) and ulcerative colitis (outcome). **(c)** MR analysis of family *Oxalobacteraceae* (exposure) and rheumatoid arthritis (outcome).

## Discussion

We report here on the relationship between host genetics and gut microbiome composition in 18,340 individuals from 24 population-based cohorts of European, Hispanic, Middle Eastern, Asian and African ancestries. We have estimated the heritability of the human gut microbiome and the effect of host genetics on the presence and abundance of individual microbial taxa. We studied the heterogeneity of the mbTL signals and characterized the impact of technical and biological factors on their effect magnitude. In addition, we explored the relevance of the identified mbTLs to health-related traits using GSEA, PheWAS and MR approaches.

Our large, multi-ethnic study allowed for an informative investigation of the human gut microbiome. However, there was large heterogeneity in the data, which reflects biological differences across the cohorts and methodological differences in the processing of samples. Overall, seven different methods of fecal DNA extraction and three different 16S rRNA regions were used^12,44^. In addition, differences in the ethnicities, ages and BMIs of the participants led to a remarkable variation in microbiome richness, diversity and composition across cohorts. Diet, medication and lifestyle, among other factors^2,3^, are known to influence the microbiome but were not included in our analysis because this data were not available for all cohorts. Large variation in the microbiome composition may have reduced the power of our mbTL analysis (see Supplementary Note).

We did not detect a host genetic effect on bacterial diversity, in line with a lack of its detectable heritability. Thirty-one taxon-specific mbTLs (20 mbQTLs and 11 mbBTLs) were identified at P<5×10^−8^. Even with our large sample size, the number of mbTLs identified is rather modest. Only the association of *LCT* locus with *Bifidobacterium* (P=1.28×10^−20^) passed the conservative study-wide significance threshold of P>1.95×10^−10^. However, we observed that heritable taxa tend to have more genome-wide significant loci and suggestively associated loci, and twin-based heritability is significantly correlated with SNP-based heritability. Our results confirm that only a subset of gut bacteria is heritable, and that the genetic architecture affecting the abundance of heritable taxa is complex and polygenic.

The association between the *LCT* locus and the *Bifidobacterium* genus was the strongest in our study. It has been shown that the functional SNP in the *LCT* locus, rs4988235, determines not only the abundance of the *Bifidobacterium* genus, but also the strength of the association between this genus and milk/dairy product intake^7^. Here, we showed the ethnic heterogeneity and age-dependent nature of the *LCT*-*Bifidobacterium* association – the effect is weaker in children and adolescents – consistent with existing knowledge on lactose intolerance^45,46^. The strongest mbQTL effect was observed in the Hispanic Community Health Study/Study of Latinos (HCHS/SOL) cohort that comprises individuals of Hispanic/Latin American ethnicity and shows the highest prevalence of the lactose intolerant NC_000002.11:g.136616754CC (rs182549) genotype (683 out of 1,097 individuals).

To explore the potential functional effects of mbTLs on health-related traits, we used GSEA, PheWAS and MR approaches. The GSEA indicated enrichment of mbQTLs for genes expressed in the small intestine and brain. These results support the existence of the gut–brain axis mediated by microbiome and likely influencing gastrointestinal, brain and mood disorders^47–49^. In addition, the PheWAS analysis identified a significant overlap between the genetic variants affecting gut microbes and a broad range of host characteristics, including psychiatric, metabolic and immunological traits, and nutritional preferences, amongst other phenotype groups (Supplementary Table 14). Moreover, genetic determinants of bacterial abundance are involved in regulating host metabolism, particularly obesity-related traits. Among the interesting bacteria, earlier studies have linked the relative abundances of *Ruminococcus*^50^, *Lachnospiraceae*^51^ and *Ruminococcaceae*^52^ to obesity. PheWAS analysis also indicated that SNPs from the *LCT* and *FUT2* loci that associated with bacterial taxa are also associated to dietary preference factors, including fish, cereal, bread, alcohol, vegetable and ground coffee intake, along with other dietary phenotypes. Interestingly, other genes found to be associated with mbTLs also included olfactory receptors (*OR1F1*) and genes involved in the absorption and metabolism of B2 and B12 vitamins (*RFK* and *CUBN*).

Genetic anchors to microbiome variation also allow for estimation of causal links with complex traits through MR approaches^53–55^. MR results indicate that Actinobacteria and *Bifidobacterium* might have a protective effect in ulcerative colitis. Cross-sectional studies have reported an increased abundance of Actinobacteria in healthy individuals as compared to inflammatory bowel disease patients^56,57^, although these results have not always been consistent^58,59^. *Bifidobacterium* was also previously shown to have a beneficial effect on ulcerative colitis in a clinical trial^58,60^. We also revealed that abundance of family *Oxalobacteraceae* in the gut microbiome might be protective for rheumatoid arthritis; the abundance of this family in lung was previously shown to be negatively associated with rheumatoid arthritis^61^. Protective effects of the bacterial taxa on these diseases support the potential of microbiome-based therapy.

In summary, we report the largest study to date to investigate the genetics of human microbiome across multiple ethnicities. Microbiome heterogeneity and high inter-individual variability substantially reduces the statistical power of microbiome-wide analyses: similar to earlier microbiome GWAS studies, we report a limited number of associated loci. Nevertheless, our results point to causal relationships between specific loci, bacterial taxa and health-related traits. Heritability estimates suggest that these associations are likely part of a larger spectrum that is undetectable in the current study sample size. This warrants future studies that should take advantage of larger sample sizes, harmonized protocols and more advanced microbiome analysis methods, including metagenomics sequencing instead of 16S profiling and quantification of bacterial cell counts. Given the essential role of the gut microbiome in the metabolism of food and drugs, our results contribute to the development of personalized nutrition and medication strategies based on both host genomics and microbiome data.

## Supporting information

Supplementary Notes, Supplementary Pictures

## Acknowledgements

We thank Jackie Senior and Kate McIntyre for editing the manuscript.

The cohorts funding and acknowledgements information is given in the Supplementary Note.

## Author contributions

A.K, A.Z., R.Kr, C.M.G., L.F. and J.R. conceived and designed the study. A.K., C.M.G., R.B., D.R. and J.W. were responsible for coordinating and performing meta-analysis. A.D., C.L.R., J.A.R.G., C.T.F., X.L., D.Z., M.J.B. lead the specific downstream analyses, and should be considered as shared second authors. Specifically, A.D. performed the PheWAS analysis, C.L.R. and C.T. F. performed the heritability analysis in TwinsUK and NTR cohorts, respectively, and J.A.R.G performed the age-related analysis of *LCT* locus. X.L. ran and interpreted the FUMA analysis, D.Z. ran and interpreted the mendelian randomization analysis. M.J.B. substantially contributed to the development of the analysis pipeline and protocols. RK, JR and AZ jointly supervised the project. A. vd G., A.C., H.J.W., Ur.V., M.J.B., S.S. and L.F. developed the pipeline for the meta-analysis and contributed to the methodology and statistical analysis. K.W. contributed to the PheWAS enrichment analysis. A.K., C.M.G., R.B., D.R., J.W., A.D., C.L.R., J.A.R.G., C.T.F., X.L., D.Z., M.J.B., M.D.A., S.S., R.Kr., J.R. and A.Z. wrote the manuscript, with contributions from all authors.

K.A.M, L.J.L and M.F collected and managed the CARDIA cohort. A.D.P, J.A.R.G., K.C., L.B. and W.T. collected and managed the GEM cohort. H.B., J.S., J.T., S.A.S, and S.J.S collected and managed the COPSAC study. D.B., O.P., T.H., T.J., and T.H.H. collected and managed the DanFunD study. D.A.H., G.F., J.R., J.W., K.H.W., M.J., N.J.T., R.Y.T., R.B. and S.V.S. collected, genotyped and managed the FGFP study. C.M.G, F.R., H.A.M., L.D. and V.W.V.J. collected and managed the Generation R study. H.N.K., H.S. and H.L.K. collected and managed the KSCS study. C.W., J.F., A.Z., L.F., S.S. and A.K. collected and managed the LLD cohort.

A.J.L., E.O., K.L., M.Lk. and M.B. collected and managed the METSIM cohort. A.A.M.M., D.M.A.E.J., D.K. and Z.M. collected and managed the MIBS-CO cohort. H.P. and ZDW collected and managed the NGRC cohort. C.T.F., D.I.B., E.J.C.G., G.E.D., G.W. and R.G.I collected and managed the NTR cohort. Da.R., E.B., E.S. and O.W. collected and managed the PNP cohort. A.A., L.A., M.D.A., Su.W. and X.L. collected and managed the PopCol cohort. A.F., C.B., M.C.R., M.Ld and W.L collected and managed the BSPSPC and FOCUS cohorts. A.G.U., C.Mv.D, Dj.R. and R.Kr. collected and managed the RS cohort. F.F., F.U.W., G.H., H.V., M.M.L, St. W. and Uw.V. collected and managed the SHIP and TREND cohorts. L.Y.M., Q.Q., R.Kn., R.C.K. and R.D.B collected and managed the SOL cohort. C.I.L.R, C.J.S., J.T.B.,

M.A.J. and T.D.S. collected and managed the TwinsUK cohort. A.A.V. and J.S.T contributed to the discussion. All authors approved the final manuscript.

## Online Methods

### Data collection

A total of 25 cohorts, comprising18,340 participants of different ethnicities and ages, participated in the microbiome GWAS analysis (Supplementary Tables 1, 2). The Supplementary Note provides detailed descriptions of data collection per cohort.

### 16S microbiome data processing

The rationale behind the selection of the 16S rRNA processing pipeline was described previously^11^. In short, the divergence in the 16S rRNA gene domains between cohorts makes operational taxonomic unit (OTU)-level analysis impossible, while the use of a direct taxonomic classification of the reads and an up-to-date reference database allowed us to achieve good between-domain concordance of taxonomic composition and a higher mapping rate.

The participating cohorts varied in their sample collection protocol, selection of DNA purification kits used to extract DNA from fecal samples, the 16S domain selected for PCR (Supplementary Table 1), read length, depth, post-sequencing quality control (QC) and the software used to merge tags of paired-end sequencing. After processing the QC-filtered merged reads, all cohorts implemented the standardized 16S processing pipeline (https://github.com/alexa-kur/miQTL_cookbook) that uses SILVA release 128^15^ as a reference database, with truncating the taxonomic resolution of the database to genus level.

Briefly, the procedure was as follows. First, all samples were rarefied to 10,000 reads using a predefined random seed to allow for rarefaction reproducibility. Samples with fewer than 10,000 reads were discarded. Second, RDP classifier v.2.12^16^ was used to bin the reads to a reference database. For each taxonomic level, the posterior probability of 0.8 was used as a cutoff to bin each read to the corresponding taxon. The posterior cutoff probability was traced for each taxonomic level separately. For example, if the posterior probability passed the cutoff on family level but not on genus level, the read was binned to taxonomy on the family level (all corresponding upper taxonomic levels) and discarded on the genus level. It was also assigned to a special “NOTAX_genus” pseudo-taxon to maintain data compositionality.

To characterize the contribution of cohort-wise metadata (16S domain, DNA extraction method, cohort ethnicity, lysis temperature and type of lysis buffer) to the microbiome composition, we used a distance-based redundancy analysis test in which each cohort represented a sample and variables represented mean abundances of genera in the corresponding cohort (taxa with prevalence below 20% discarded). The association of metadata with richness was performed by multivariate linear regression analysis.

The alpha diversity indices, including Shannon, Simpson and inverse Simpson indices, were calculated on genus level with non-adjusted, non-transformed taxa counts. For all other analyses, the taxonomic counts of non-zero samples were natural log–transformed and adjusted for potential covariate effects using linear regression. The list of covariates used in the regression models varied between cohorts, but always included sex, age, genetic principal components (PCs) calculated on non-imputed genetic data (3 PCs for monoethnic cohorts, 10 PCs for multiethnic cohorts and 5 PCs for the HCHS/SOL cohort as a multi-ethnic population of different, but closely related ethnicities; see Supplementary Note for Cohort descriptions) and cohort-specific potential microbiome batch effects, if applicable. Variables such as the length of time in non-frozen storage, the 16S sequencing batch, etc. were also included. The residuals of the adjustment were then scaled and centered (mean=0 and SD=1).

In the analysis of microbiome composition heterogeneity, the cohorts SHIP/SHIP-TREND and GEM_HCE_v12/GEM_HCE_v24/GEM_HCE_ICHIP were merged to SHIP and GEM, respectively, because they were analyzed with exactly the same protocols in the same laboratories. In the microbiome–genetics analysis, these five cohorts were included individually as they differed in the genotyping arrays and/or general populations they represented.

For each cohort, only the taxa present in more than 10% of the samples were included in the quantitative microbiome trait loci (mbQTL) mapping, whereas taxa present in more than 10% but less than 90% of the samples were included in binary trait loci (mbBTL) mapping (Supplementary Table 3). Study-wide cutoffs for mbQTL mapping included an effective sample size of at least 3,000 samples and presence in at least three cohorts. For mbBTLs, a mean abundance higher than 1% in the taxon-positive samples was required. This resulted in 211 taxa (131 genera, 35 families, 20 orders, 16 classes and 9 phyla) that passed taxon inclusion cutoffs for mbQTL analysis, and 177 taxa (108 genera, 34 families, 16 orders, 12 classes and 7 phyla) for mbBTL analysis.

### Genetic data processing

Despite the difference in genotyping array platforms, most cohorts used similar procedures for imputation and post-imputation filtering steps. Twenty-three out of 24 cohorts used the Michigan Imputation Server (https://imputationserver.sph.umich.edu/index.html) for imputation, using the HRC 1.0 or 1.1 reference panel^62^. Due to restrictions in manipulating data, the PNP study employed an in-house pipeline for imputation instead, using IMPUTE2^63,64^ software (v.2.3.2) and 1000G reference panel with addition of population-matched genotypes of Jewish individuals^65^. The post-imputation cutoffs were the same for PNP and the other cohorts.

Post-imputation VCFs were transformed into TriTyper format and filtered using GenotypeHarmonizer v.1.4.20 software^66^. The following cutoffs were applied for inclusion: minor allele frequency >0.05, pointwise imputation QC >0.4 and SNP-wise call rate filtering >0.95.

### Heritability analysis

Heritability was calculated using data collected on 169 MZ and 419 DZ pairs of twins from the TwinsUK cohort (total of 1,176 individuals). Twin-based heritability was calculated by fitting an ACE model using the OpenMx package (v.2.8.3), as previously described^5^. Prior to heritability estimation, the taxonomic abundance was normalized using inverse rank sum transformation.

Since the NTR cohort comprised only MZ twins, the between-cohort heritability concordance was calculated as the correlation of intraclass correlation coefficient (ICC) for MZ twins. Pearson’s correlation between NTR’s and TwinsUK’s ICCs was used to estimate the concordance. For mbQTLs, SNP-based heritability was calculated by LD score regression using ‘LDSC’ tool^67^.

### Microbiome GWAS analysis

The modified version of the eQTL mapping pipeline (https://github.com/molgenis/systemsgenetics/tree/master/eqtl-mapping-pipeline) was used to perform mbQTL mapping^68^.

The microbiome GWAS was performed in three ways. First, we performed GWAS on three microbiome alpha diversity metrics (Shannon, Simpson and Inverse Simpson), using Spearman correlation between SNP dosages and alpha diversity metrics after adjustment for age, sex, technical covariates and genetic principal components.

Second, we used Spearman correlation to identify loci that affect the covariate-adjusted abundance of bacterial taxa, excluding samples with zero abundance (mbQTLs).

Third, we identified the loci associated with probability of presence vs absence of the bacterial taxon (mbBTLs). To perform mbBTL analysis, we used a two-stage approach composed of fast correlation screening followed by logistic regression analysis as a robust method for binary traits GWAS^19^. First, we calculated the Pearson correlation between SNP dosage and bacterial presence encoded as 0/1, without adjusting for any covariate effect and using the previously mentioned eQTL mapping pipeline, and used weighted Z-score meta-analysis to calculate non-centrality for SNP-taxon association. Finally, all SNP-taxon pairs with a first stage meta P-value <1×10^−4^ were recalculated using multiple logistic regression (R base package, versions from 3.2.0 to 3.5.1 depending on the group) with bacterial presence as an outcome and using SNP dosage along with the list of covariates as predictors. All the mbBTLs that reached nominal genome-wide significance threshold (P<5×10^−8^) in logistic regression analysis had a Pearson correlation P-value (at first stage) more significant than P<10^−6^, presuming the completeness of two-stage procedure in revealing genome-wide significant mbBTL using P<10^−4^ cutoff at the first stage of analysis.

### mbTL meta-analysis

Meta-analysis was performed using a weighted Z-score method implemented in BinaryMetaAnalyzer (v.1.0.13B available on MiBioGen Cookbook), a part of the eQTL mapping pipeline that was used in large-scale eQTL meta-analyses^20,68^. Per-cohort, Z-scores were calculated from Spearman correlation p-values using inverse normal transformation, transforming two-tailed p-values to one-tailed p-values and tracing the effect directions using the following formula:

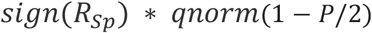

Where *sign*(*R*_*SP*_) denotes the sign of Spearman correlation, *qnorm* denotes the quantile function for the normal distribution and *P* denotes the two-tailed p-value of Spearman correlation. For quantitative mbQTLs, the cohorts were weighted by the square root of the effective sample size (the number of samples having the bacterial taxon). For binary mbQTLs, the square root of the reported cohort size was used as a weighting for each study. The summary statistics generated for mbQTLs also include meta-effect sizes and standard errors. These were generated using the inverse variance weighted meta-analysis method performed on the per-cohort effect sizes and standard errors, backtracked from association Z-scores and minor allele frequencies using the strategy proposed and implemented by Zhu *et al*^69^, where they also give the detailed derivation of the following equations:

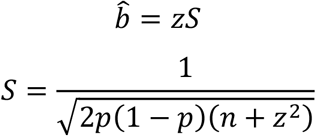

Where, *b* is the estimated effect size, *S* is the estimated standard error, *p* is the allele frequency and *n* is the sample size.

### Heterogeneity exploration analysis

Cross-study heterogeneity of the effects of genetic variants in the relative abundance of taxonomical units was assessed using Cochran’s Q-test for heterogeneity^70^, as implemented in METAL v2018-08-28^71^, for all genome-wide significant variants (P<5×10^−8^) found in our main analysis. To avoid reporting false-positive associations due to different study designs or data collection methods, we used a stringent threshold of P<0.05 to reject the null hypothesis of no heterogeneity. This threshold is conservative considering that several variants were tested simultaneously, and no correction for multiple testing was applied. When there was evidence of heterogeneity, a random effect model was also implemented at the meta-analysis level to confirm the association results, using the metaphor R package v.2.0-0 (https://cran.r-project.org/web/packages/metafor/metafor.pdf).

Additionally, when there was evidence for heterogeneity of a SNP-effect across cohorts, we implemented a meta-regression approach using the same package to assess whether variables such as age, ethnicity or sequenced region could explain the observed effect-size heterogeneity.

### Analysis of SNP–age interaction analysis in the *LCT* locus

To discover whether the association of functional SNPs in the *LCT* locus to the abundance of the *Bifidobacterium* genus varied between groups of adults and infants, we performed age–SNP interaction analysis in the GEM cohort, which comprises three sub-cohorts that each have a comparable number of individuals above and below puberty age. The age of 17 years was selected to split the cohort into the age groups: adolescents or adults. Since the GEM cohort was composed of three sub-cohorts of different ethnic composition, we evaluated the interaction in both joint analysis and in each subcohort separately, using the following formula:

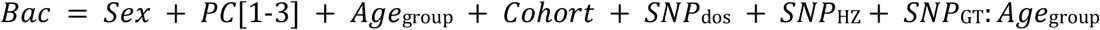

where *Bac* is the log-transformed count of genus Bifidobacterium, *PC*[1-3] are three floats with the first 3 genetic PCs, *Cohort* is a batch variable that determines the cohort the sample belongs to, *SNP*_*dos*_ is a float-encoded dosage of alternative allele, *SNP*_*HZ*_ is a Boolean variable describing heterozygosity, *SNP*_*GT*_ is a genotype encoded as an unordered factor and *Age*_*group*_ is a two-level factor (above or below split level). The inclusion of a numeric dosage variable and a Boolean SNP_HZ_ variable allowed us to properly adjust for the recessive effect of the SNP on *Bifidobacterium* abundance without neglecting SNP imputation uncertainty as embedded in SNP dosage.

The analysis was then repeated for each GEM subcohort separately, using the same model.

### Association of mbTL-associated taxa with host phenotypes

Bacterial taxa found to be significantly associated with genetic determinants were correlated with 207 host phenotypes, including the intrinsic host properties, diet, disease and medication information, in the LLD and FGFP cohorts. We used Spearman correlation with Benjamini-Hochberg (BH)-adjustment for multiple testing to assess the correlation between phenotypes and bacteria that had mbQTLs. For the taxa with mbQTLs, samples with zero abundance were truncated. For the taxa with mbBTLs, the abundance was transformed to a binary trait encoding presence/absence.

### FUMA analyses of meta-analysis results

Functional mapping and annotation of 30 meta-analysis results were performed with FUMA (v1.3.5), an integrated web-based platform^31^. Summary statistics from the mbQTL analyses for each of the 20 independent association signals were used in the analysis. Genome-wide significant loci and their boundaries were defined as non-overlapping genomic regions that extend across an LD window of r^2^≥0.4 (based on the 1000G EUR reference panel)^72^ from the association signals with P<5.0×10^−8^. Independent (r^2^<0.1) lead SNPs from each locus were defined as those most strongly associated with a microbial trait (i.e. with the lowest P value) at the specific region. Multiple risk loci were merged into a single genomic locus if the distance between their LD blocks was <250 kb.

Functional annotation of all candidate risk SNPs was obtained from different repositories integrated in FUMA. Furthermore, these functionally annotated SNPs were mapped to protein-coding genes using the following two strategies: (1) positional mapping, with the maximum distance of 10 kb to protein-coding genes, and (2) eQTL mapping, using information from data repositories such as GTEx v7 and Blood eQTL browser (http://genenetwork.nl/bloodeqtlbrowser/)^20^.

As the mbBTL mapping procedure provides accurate statistics for only the subset of SNPs (see Microbiome GWAS analyses paragraph), and we thus lack full summary statistics, we only performed positional mapping for mbBTLs, taking in the protein-coding genes within 10 kb distance of the 10 leading SNPs per trait.

All mapped protein-coding genes were combined into one list for either mbQTL or mbBTL analysis prior to performing GSEA integrated in FUMA. In further investigations, hypergeometric tests of enrichment of all mapped genes were performed not only in tissue-specific (differentially expressed) gene sets, but also in gene sets curated from various sources, e.g. MsigDB. We reported all enriched gene sets (≥2) with an FDR adjusted P-value <0.05.

### PheWAS, genetic correlation and colocalization analysis

We performed the PheWAS look-ups in the summary statistics results of 4,155 traits collected by the GWASATLAS^32^ (http://atlas.ctglab.nl/, accessed on: 25-09-2019) database for the top SNPs per mbQTL locus that were revealed by either mbQTL or mbBTL mapping. GWASATLAS includes 600 traits from the UK Biobank and is enriched with extensive phenotypes on proteomics (n=1124 proteins), hematology (n=36), metabolomics (n=1145 metabolic features) and immune markers (n=241), studied across variable sample sizes. It also contains 1,009 GWASs performed prior to the UK Biobank effort, all categorized under 27 phenotype domains. Next, we tested if any of these 27 domains were enriched by the phenotypes associated to one of the SNPs of interest (using a liberal P-value threshold of 0.05 for the SNP–phenotype association) as compared to the expected distributions under the null hypothesis. In order to obtain the distributions under the null hypothesis, we selected matching 1000 SNPs for each top SNP using SNPSNAP^73^ matched by allele frequency, gene density, number of LD pairs and distance from the closest gene.

We then extracted corresponding results from the GWASATLAS for the matched 30,000 SNPs (1000 matching SNPs per each top mbTL SNP). The enrichment of each domain was tested by comparing the proportions of observed and expected significant results for the SNPs of interest using the prop.test function in R. This resulted in one-sided P-values and odds ratios. Seven domains (Aging, Body structures, Connective tissue, Ear-Nose-Throat, Infection, Muscular and Social Interactions) that included fewer than 20 GWAS tables were excluded from the enrichment tests, resulting in 20 domains. We used a conservative Bonferroni-based P-value threshold of 8.06×10^−5^ for the enrichment testing, accounting for 20 domains and a total of 30 mbTL top SNPs coming from both the mbQTL and mbBTL mapping. In addition, we performed gene-based PheWAS look-ups in the GWASATLAS for candidate genes of interest within 250 kb around the association peaks, as defined by the FUMA algorithms.

The genetic correlation between Bifidobacterium and its PheWAS-related traits (from Table S12) was estimated following a LD-score regression approach^67^ using the ‘ldsc’ tool. For testing colocalization of the PheWAS signals, we used the approximate Bayes factor approach as implemented by the “coloc.abf” function from the “coloc” library in R^74^, using genetic variants within ±250 kb around the top signals.

### Mendelian Randomization analysis

MR analyses were performed in R using TwoSampleMR package (v.0.5.5)^36^. Causality direction was tested between the microbiome and two data types: (1) autoimmune, cardiovascular, metabolic (including weight-related phenotypes) and psychological diseases (GWAS summary statistics from MRBase^36^) known to be associated with microbiome composition^2,3,37–42,47^ and (2) 42 nutritional phenotypes and alcohol intake frequency from the UK Biobank round 2 (http://www.nealelab.is/uk-biobank/).

For MR analyses, the combined meta-effects and standard errors from inverse variance meta-analysis were used.

To test if a complex trait affected microbiome composition, we selected independent genetic variants associated with complex traits at the genome-wide significant level (P<5×10^−8^) and used these as instruments in our MR analyses. For complex diseases, we transformed Odd Ratios (ORs) and C.I. to effect sizes and standard errors using the built-in function of the TwoSampleMR package. To test if microbiome changes were causally linked to complex traits, we first confined ourselves to bacteria with genome-wide significant QTLs. For these, we selected all SNPs with a less stringent cut-off of P<1×10^−5^ in our MR analyses as instruments. This strategy was used to increase the number of SNPs available in order to perform sensitivity analyses, as shown previously^53^. Independent SNPs were selected as instrumental variables based on r^2^ < 0.001 in 1000G EUR data, within the TwoSampleMR package. When no shared SNPs were available between exposure and outcome, proxies from the 1000G EUR data (r^2^ > 0.8) were added. We kept only the results based on at least three shared SNPs. MR causality tests were performed using the Wald ratio, and Wald ratios were meta-analyzed using the inverse-variance weighted (IVW) method^75^. We also estimated the causality using additional methods: the weighted mode method^76^, which provides an alternative approach to IVW; MR-Egger^77^, which estimates the degree of horizontal pleiotropy in the data; and MR PRESSO^78^, which estimates the pleiotropy and corrects for it by removing outliers from the IVW model. We also assessed the heterogeneity of the results using Cochran’s Q statistics^75^ and using leave-one-out analyses^36^. We estimated instrument variable (IV) strengths using F statistics: the amount of variance explained by IVs was calculated for each exposure using the TwoSampleMR package (get_r_from_lor function) for binary traits and VPE as defined in Shi *et a*l^79^. F statistics were then calculated as 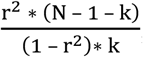, where r^2^ is the variance explained, N is the sample size and k is the number of IVs. We kept the results for the conventional threshold of F statistics >10 ^80^.

After performing the MR tests, we excluded duplicated GWAS traits, as the same phenotype is often studied in multiple GWAS. To remove the duplicates, we kept the study with the largest sample size among all the tested GWAS studies for each trait.

After excluding duplicates and tests performed with weak instruments (F statistics <10), we applied a BH correction for multiple testing to the results obtained from the IVW MR test, and subsequently used a stringent filtering procedure on the significant results to avoid false-positives. Specifically, we removed the MR results that were based on fewer than three SNPs and thus could not be further investigated with sensitivity analyses. We also removed the MR results that were not supported by other MR tests (weighted mode method P >0.05, MR PRESSO P>0.05) and those that showed substantial pleiotropy or heterogeneity as estimated by MR-Egger (MR-Egger intercept P<0.05) or MR PRESSO outliers-adjusted test (P>0.05), as well as those where leave-one-out analysis identified one SNP driving the signal (all but one leave-one-out configurations had P<0.05). Of note, MR-Egger slope, which represents the causal estimate, was not used as a filtering step given the reduced power to detect causal effects. It is also worth noting that for all but one of the reported MR results that passed all the filters above, the MR-Egger slope p-value was greater than 0.05, therefore an MR-Egger intercept P<0.05 cannot be used to exclude presence of pleiotropy. Even though many of our MR-Egger intercept results provided little evidence of directional pleiotropy, it is worth noting that a P<0.05 cannot exclude the presence of pleiotropy and requires further understanding of the biological mechanisms underpinning the relationship between genetic variation, the gut microbiome and health outcomes. To exclude more complex causality scenarios, we also removed those results for which the reverse MR P-value was below 0.05. Of note, the causal relationship identified for the microbiome feature class Actinobacteria (as exposure) and ulcerative colitis (outcome) showed a consistent effect direction when using only the only genome-wide significant SNP, but with wider confidence interval (OR=0.40 [95% CI: 0.22-0.71] P_nominal_=0.002).

## Data availability statement

Full GWAS summary statistics for mbQTLs are available at www.mibiogen.org website built using the MOLGENIS framework^81^.

16S data availability:

BSPSPC and FOCUS data is available from Sequence Read Archive (SRA), PRJNA673102 All CARDIA data, including 16S rRNA sequencing, cannot be made available on publicly available databases due to the confidentiality restrictions. The data can be requested from CARDIA Study Data Coordinating Center at the University of Alabama at Birmingham, following CARDIA Confidentiality Certification rules. The process for obtaining data through CARDIA is outlined at: https://www.cardia.dopm.uab.edu/publications-2/publications-documents.

COPSAC data is available on SRA (PRJNA683912).

DanFunD is not deposited on the public databases due to the legal and ethical restrictions. Access to the data and biological material can be granted by the DanFunD steering committee (https://www.frederiksberghospital.dk/ckff/sektioner/SBE/danfund/Sider/How-to-collaborate.aspx).

FGFP data is available on European Genome-Phenome Archive (EGA), EGAS00001004420 GEM data is available on SRA (PRJEB14839).

Generation R and Rotterdam Study data cannot be made publicly available due to ethical and legal restrictions; these data are available upon request to the data manager of the Rotterdam Study Frank van Rooij or of the Generation R Study Claudia Kruithof and subject to local rules and regulations.

HCHS/SOL data is available from ENA (European Nucleotide Archive), ERP117287.

KSCS data is available at the public repository, Clinical and Omics data archives (CODA) in the Korea National Institute of Health by accession number R000635 (http://coda.nih.go.kr/coda/coda/search/omics/genome/selectSearchOmicsGenomePop/R000635.do).

LLD and MIBS data are available from EGA, EGAS00001001704, EGAS0000100924). METSIM data is available on SRA (SRP097785).

NGRC data is available on ENA (ERP016332).

NTR has a data access committee that reviews data requests and will make data available to interested researchers. The data come from extended twin families and pedigree structures with twins, which create privacy concerns and thus cannot be shared on publicly available databases. Researchers may contact prof Eco de Geus for data request..

PNP is available on ENA (PRJEB11532).

POPCOL is available on EGA (EGAS00001004869).

SHIP and SHIP-TREND data can be obtained from the SHIP data management unit and can be applied for online through a data access application form (https://www.fvcm.med.uni-greifswald.de/dd_service/data_use_intro.php)

TwinsUK data is available on the European Nucleotide Archive (ENA, accession ERP015317).

## Code availability statement

All code used in the study is available on the Consortium GitHub (https://github.com/alexa-kur/miQTL_cookbook) or on the websites of corresponding software packages.

