## Supplementary Notes, Supplementary Pictures for "Large-scale association analyses identify host factors influencing human gut microbiome composition"

### Supplementary Notes and Figures

#### Supplementary Note 1. Cohort Descriptions

<u>BSPSPC (PopGen)</u>

The PopGen cohort (mean age 61.5 (16.6), 55% male) is a population-based cohort from the area around Kiel, Schleswig-Holstein, Germany. Participants were genotyped using the Affymetrix Genome-Wide Human SNP Array 6.0. Fecal samples of 714 individuals were collected by the participants themselves at home in standard fecal collection tubes and shipped to the study center, where they were stored at -80°C until processing. DNA from fecal samples (app. 200 mg) was extracted using the QIAamp DNA stool mini kit automated on the QIAcube. Genotyping data generation, extraction of fecal DNA and sequencing of the V1-V2 variable region of the 16S rRNA gene and all data processing were performed at the Institute of Clinical Molecular Biology, Kiel, Germany. The study was approved by the institutional ethical review committee of Kiel University, Germany. Written informed consent was obtained from all participants.

<u>CARDIA (Coronary Artery Risk Development in Young Adults Study)</u>

Coronary Artery Risk Development in Young Adults Study (CARDIA) is a population-based prospective study of the evolution of cardiometabolic disease. African American and European American adults were recruited from four U.S. urban areas (Birmingham, AL; Chicago, IL; Minneapolis, MN; Oakland, CA in 1985-1986) (n=5,115, aged 18-30). They have subsequently been examined nine times. A microbiome study was initiated at the Year 30 follow-up examination (2015-2016) in a subset of participants (n=615) who had not taken antibiotics in the past month. Fecal DNA was extracted with the MoBio PowerSoil kit, and the V3-V4 region of the 16S rRNA gene was sequenced with Illumina MiSeq (2x300bp) at HudsonAlpha Institute for Biotechnology (Huntsville, AL, USA). A subset of cohort participants has been genotyped with

the Affymetrix Genome-Wide Human SNP Array 6.0. After quality control and removal of participants with non-overlapping data on microbiome and host genetics, data from 114 African Americans and 257 European Americans (total n=371) were available for analysis.
The study was approved by Institutional Review Boards of University of Alabama at Birmingham, Birmingham, AL, Kaiser Permanente Division of Research, Oakland CA, University of Minnesota, Minneapolis, MN, and Northwestern University, Chicago, IL. Written informed consent was obtained from all participants.

COPSAC$_{2010}$

The Copenhagen Prospective Studies on Asthma in Childhood 2010 (COPSAC$_{2010}$) cohort is a prospective mother-child cohort of 700 children and their families, recruited during week 24 of pregnancy, with written informed consent obtained from all mothers[82]. The participants reside in and around Copenhagen, Denmark. The design builds upon the previous COPSAC$_{2000}$ cohort[83] and is based on detailed longitudinal clinical assessments of asthma, allergy, eczema and other outcomes. Blood tests were taken from the infants at age of six months, and DNA was extracted from plasma. Genome-wide genotyping was performed using the Illumina OmniExpress-8 v1.4 and Exome BeadChip. Fecal samples were collected at visits to the clinic or at home by parents using detailed instructions at ages 1 week, 1 month, and 1, 4, 5, and 6 years. For the present study, samples for age 4-6 years were used. Genomic DNA was extracted from the child's samples using the PowerMag® Soil DNA Isolation Kit, and the V4 region of the 16S rRNA gene was amplified and sequenced on an Illumina MiSeq system, as previously described in detail[84]. At the relevant timepoint, we had both genotype and microbiome data for 380 children to include in this study, 73 of whom had taken antibiotics in the six months before the fecal sample date. The study was approved by Danish Ethics Committee (H-B-2008-093) and the Danish Data Protection Agency (2008-41-2599).

DanFunD (The Danish study of Functional Disorders)

DanFunD is a population-based cohort initiated to outline the epidemiology of functional somatic syndromes[85]. The study population comprises a random sample of 9,656 men and women aged 18-76 years from the general population who were examined from 2011 to 2015. Genotyping using the Human OmniExpress Bead Array (Illumina Inc., San Diego, CA, USA) was conducted on human leukocyte DNA for the entire cohort. A subset of 2,464 participants volunteered to provide a fecal sample collected under standardized conditions. Microbial DNA extraction using the NucleoSpin Soil kit (Macherey-Nagel, Düren, Germany) and subsequent sequencing of the hypervariable region V4 of the bacterial 16S rRNA gene on an Illumina HiSeq 2500 platform was conducted at Beijing Genomics Institute (BGI Europe, Copenhagen, Denmark). In total, 2,396 samples passed the QC for genotyping and 16S sequencing and were included in the GWAS.

The study was approved by Ethical Committee of Copenhagen County (Ethics Committee: KA-2006-0011; H-3-2011-081; H-3-2012-0015) and the Danish Data Protection Agency. Written informed consent was obtained from all participants.

FGFP (Flemish Gut Flora Project)

The FGFP is a population-based study cohort of 2,482 individuals from the Flanders region of Belgium. Blood and stool samples of volunteers were collected between June 2013 and April 2016. Genotyping was performed using the Human Core Exome arrays v1.0 and v1.1. Sampling kits were sent to the volunteer's homes and stored there at -18°C until collection and storage in the Raes Lab facilities at -80°C. DNA was extracted from the frozen fecal samples using the PowerMicrobiome RNA Isolation Kit, as described in Falony et al[3]. Sequencing of the V4 region of the16S rRNA gene was carried out on the Illumina HiSeq platform. After quality control, 2,259 samples had genotype and 16S data (1,328 females, 896 males, mean age 52.3 yrs). FGFP procedures were approved by the medical ethics committee of the University of Brussels–Brussels University Hospital (approval 143201215505, 5/12/2012). A declaration concerning the FGFP's privacy policy was submitted to the Belgian Commission for the Protection of Privacy. Written informed consent was obtained from all participants.

FOCUS

The FoCus cohort (mean age 51.4(14.6) yrs, 42% male) is a population-based cohort from the area around Kiel, Schleswig-Holstein, Germany, and part of the competence network Food Chain Plus (FoCus, http://www.focus.uni-kiel.de/component/content/article/88.html). Participants were genotyped using the Infinium OmniExpressExome Array. All data generation and processing was performed at the Institute of Clinical Molecular Biology, Kiel, Germany, similar to the PopGen cohort.

The study was approved by the institutional ethical review committee of Kiel University. Written informed consent was obtained from all participants.

GEM (The CCC GEM project)

The CCC GEM project is a prospective international research study designed to identify the potential triggers that contribute to the onset of Crohn's Disease. Since 2008, the GEM project has recruited over 5,000 healthy first-degree relatives of Crohn's Disease patients with an age range of 6-35 years. At the time of recruitment, participants were screened using a standardized questionnaire to exclude any history or symptoms of inflammatory bowel disease or other gastrointestinal diseases. For the microbiome GWAS, we used data from participants recruited in Canada (n=1,115), the United States (n=17) and Israel (n=111). Stool DNA was extracted using the QIAamp DNA Stool Mini Kit (Qiagen, Hilden, Germany). The V4 hypervariable region of bacterial 16S ribosomal RNA (16S rRNA) was sequenced using a MiSeq platform (Illumina Inc. San Diego, CA, USA) and primers 515F/806R[86]. Genotyping of the cohort was performed using the HumanCoreEXOME-12v1.1 chip (n=379), HumanCoreEXOME-24v1.0 chip (n=203) and both ImmunoChip and HumanCoreEXOME-12v1.1 chip (n=662) (Illumina, Inc. San Diego, CA, USA). Thus, in mbQTL mapping, the cohort was split into subcohorts GEM_v12, GEM_v24 and GEM_ICHIP respectively. Among subcohorts, GEM_v24 mostly comprises individuals of Israeli ethnicity (70%, 61 Ashkenazi, 34 Sephardic, 18 other/unknown subethnicities), while the other two subcohorts are of a European ancestry. Only one sample from one member from each family enrolled in the project was included in the current microbiome GWAS study. After stringent QC, as previously described[9], the overlap between samples with genotyping and microbial 16S sequencing data yielded 1,243 samples (676 females, 567 males, median age 19.0(8.03) yrs) for use in the microbiome GWAS analysis. None had used antibiotics in the three months before fecal collection.

The study was approved by Mount Sinai Hospital Research Ethics Board (Toronto–Managing Center) and local centers. Written informed consent was obtained from all participants.

GenR (The Generation R Study)

GenR is a population-based, prospective, multi-ethnic pregnancy cohort study from fetal life until young adulthood. It is conducted in the city of Rotterdam, the Netherlands[87]. Genotyping of this cohort was performed using Illumina HumanHap 610K[88]. Stool sample collection started in

2012 and comprised 2,111 children. Fecal DNA was extracted using DiaSorin Arrow DNA (Isogen Life Science, De Meern, the Netherlands) with a bead-beating step. Sequencing of bacterial 16S gene, domain V3-V4, was performed in the Laboratory of Human Genetics at Erasmus MC Rotterdam using the Illumina MiSeq platform[89]. After stringent QC, the overlap between samples with genotyping and microbial 16S sequencing data yielded 1,328 samples (656 females, 672 males, mean age 9.8(0.3) years) for use in the microbiome GWAS analysis. None had used antibiotics in the six months before fecal collection.

The study was approved by the Medical Ethical Committee of Erasmus MC, University Medical Center Rotterdam. Written informed consent was obtained from all participants.

KSCS (Kangbuk Samsung Cohort Study)

The KSCS is a prospective cohort study to evaluate the natural history, prognosis and genetic and environmental determinants of a wide range of health traits and diseases among Korean adults. There are two major cohort studies in Kangbuk Samsung Hospital: the KSCS and the Kangbuk Samsung Health Study (KSHS). The KSHS is a retrospective cohort study using de-identified data routinely collected during health screening visits from 2002 to present, and includes standardized and high quality clinical, imaging and laboratory procedures and information on multiple lifestyle and medical conditions. The KSCS is a prospective study that has now started to apply more strict and standardized procedures. They obtained informed consent for data linkage to national registries for death, cancer and medical utilization since 2011. Genotyping was conducted using the Illumina HumanCore BeadChips 12v in 2014 (n=2,040). Fecal samples were collected from 1,463 participants between June and September 2014. DNA extraction from fecal samples was performed within one month of storage using the MoBio PowerSoil ® DNA Isolation Kit (MO BIO Laboratories, Carlsbad, CA, USA). Sequencing of the bacterial 16S rRNA gene, domain V3-V4, was performed using the Illumina MiSeq platform (Illumina, San Diego, CA, USA). After QC, 811 samples (319 females, 492 males, mean age 44.1 yrs) with overlapping genotype and 16S data were included in the microbiome GWAS.

The study was approved by the EUMC review board 2014-06-024 and KBSMC review board 2013-01-245. Written informed consent was obtained from all participants.

LLD (LifeLines-DEEP)

LLD is a subcohort of the prospective LifeLines cohort from the northern provinces of the Netherlands (Groningen, Drenthe and Friesland) and includes participant of Dutch ethnicity. Blood and fecal samples of LLD participants were collected between April and August 2013. Genotyping was performed using the Illumina ImmunoChip and Illumina Human CytoSNP-12 microarrays. Fecal DNA was extracted using the Qiagen AllPrep kit with a bead-beating step. Sequencing of the bacterial 16S gene, domain V4, was performed at the Broad Institute (Boston, USA) using the Illumina MiSeq platform. The overlap between samples with genotyping and microbial 16S sequencing data yielded 875 samples (504 females, 371 males, mean age 45.4(13.3) yrs) used for the microbiome GWAS analysis. Of these, 70 participants were PPI users and eight people used antibiotics in the six months prior to fecal collection. Each participant signed an informed consent form before participation in the cohort according to the UMCG Institutional Review Board (IRB; #M12.113965).

METSIM (METabolic Syndrome In Men)

The METSIM cohort is a longitudinal population-based cross-sectional cohort comprising 10,197 randomly selected non-diabetic Finnish men (aged 45 to 73 years) who were examined in 2005-2010. Genotyping was performed using the Illumina Omni ExpressExome microarray. Microbial DNA was extracted from frozen fecal samples using the PowerSoil DNA Isolation Kit (MO BIO Laboratories, Carlsbad, CA, USA) following the manufacturer's instructions. Amplification of the V4 hypervariable region of the 16S rRNA gene was done using the 515F and 806R primer and sequenced with the Illumina MiSeq platform at the University of California, Los Angeles. For the current microbiome GWAS study, we used a subset of the METSIM cohort consisting of 522 samples (mean age 61.91 (5.42) yrs) with overlapping genotyping and microbial 16S sequencing data. The study was approved by Ethics Committee of the Northern Savo Hospital District, Finland. Written informed consent was obtained from all participants.

MIBS (Maastricht Irritable Bowel Syndrome)

The MIBS cohort with biobank aims to identify subgroups of IBS according to phenotypical and genotypical characterization. At present, it includes 520 subjects with a clinical diagnosis of IBS according to the Rome III criteria (from primary-tertiary care) and 220 age- and gender-matched healthy controls. At baseline, all subjects completed an extensive questionnaire on demographics, lifestyle factors, medical history and medication use, as well as a 14-day symptom dairy, the GSRS, HADS, STAI, SF-36 and a food frequency questionnaire. In addition, blood (serum, (platelet-poor) plasma, DNA), feces and exhaled air were collected. In subgroups, a rectal barostat and multisugar test for intestinal permeability was performed. All participants gave written informed consent. For the present microbiome GWAS study, only controls (N=80, mean age 48.7(18.2), 43% male) were included. Genotyping was performed using the Illumina ImmunoChip and Illumina Human CytoSNP-12 microarrays. Fecal DNA was extracted using the Qiagen AllPrep kit with bead-beating step. Sequencing of bacterial 16S gene, domain V4, was performed at the Broad Institute (Boston, USA) using the Illumina MiSeq platform. Each participant signed an informed consent form before participation in the cohort according to the Maastricht University Medical Center (MUMC+) IRB (#MEC 08-2.066.7/pl).

NGRC (NeurGenetics Research Consortium)

The NGRC is a collaborative study of gene-environment-microbiome interaction on Parkinson's disease. It is being conducted in the United States. A GWAS was conducted in 2009 on DNA from whole blood on the Illumina HumanOmni1-Quad_v1-0_B array[90]. Stool and metadata were collected from a subset of participants in 2014. Fecal DNA was extracted using the MoBio PowerMag Soil DNA Isolation Kit (Optimized for KingFisher). The 16S rRNA V4 amplicon was sequenced using the Illumina MiSeq platform[91]. For the microbiome GWAS study, only 133 control participants were used. These controls were free of neurodegenerative disease, had a mean age of 71.9 (7.5) years old and 58% were female.

The study was approved by the institutional review boards of the participating institutions: Albany Medical Center, Emory University; Kaiser Permanente Northwest Division, New York State Department of Health, Oregon Health & Sciences University (OHSU) and the Department of Veterans Affairs VA Puget Sound Health Care System (VAPSHCS). Written informed consent was obtained from all participants.

NTR (the Netherlands Twin Registry)

The NTR collects data and biological samples on Dutch multiples and their family members[92]. The NTR samples included in the microbiome GWAS were collected for two separate studies: the first focused on the association between obesity and the gut microbiome and the other collected samples from family members and spouses. Genotyping was performed on the

Affymetrix SNP 6.0, Affymetrix Axiom and Illumina GSA arrays. Fecal DNA was extracted using the Qiagen PowerSoil kit with the addition of the heating step of the Qiagen PowerFecal kit. The sequencing of the V4 domain of the 16S gene was performed using the Illumina MiSeq platform. DNA extractions and sequencing were performed at the Avera Institute for Human Genetics (Sioux Falls, SD, USA). One of each twin pair was randomly selected for inclusion in the GWAS analyses (156 twin pairs, 123 unrelated individuals, 279 individuals total, mean age 35.4(12), 29.8% male). Both MZ twins were included for the ICC calculations between MZ twin pairs for comparison with heritability estimates (156 twin pairs). None of the participants reported using antibiotics within six months of fecal collection.

The study was approved by Central Ethics Committee on Research involving human subjects of the VU University Medical Center, Amsterdam. Written informed consent was obtained from all participants.

PNP (Personalized Nutrition Project)

The PNP is a large-scale nutrition initiative in Israel that aims to help people make food choices that would normalize their blood glucose level and improve their health and well-being. The cohort has over 1,000 healthy individuals of Israeli ethnicity living in Israel and aged between 18 and 70 years. The cohort consists of self-reported Ashkenazi (n=508), North African (n=64), Middle Eastern (n=34), Sephardi (n=19), Yemenite (n=13) and 'admixed/other' (n=408) ancestries. The top two host genetic principal components (PCs) are strongly associated with self-reported ancestry ($P<10^{-32}$ for both PC1 and PC2, Kruskal-Wallis test). Participants were genotyped using Illumina OMNI-EXPRESS arrays. They also provided stool samples, which were collected using a swab or an OMNIGENE-GUT (OMR-200; DNA Genotek) stool collection kit. Metagenomic sequencing was performed on DNA extracted from the stool samples as was 16S rRNA profiling by sequencing the V3-V4 region. 481 individuals were included in the current study (mean age 43.7(13.1), 36.4% male).

The study was approved by Tel Aviv Sourasky Medical Center Institutional Review Board (IRB), approval numbers TLV-0658-12, TLV-0050-13 and TLV-0522-10; Kfar Shaul Hospital IRB, approval number 0-73; and Weizmann Institute of Science Bioethics and Embryonic Stem Cell Research oversight committee. Written informed consent was obtained from all participants.

PopCol (Population-based Colonoscopy)

PopCol is a cohort study in Stockholm, Sweden that includes a data-rich set of individuals with data available from bowel symptoms questionnaires, gastroenterology visits and biospecimens (genotype and 16S sequencing from blood and stool samples, respectively)[93,94]. Genotyping was carried out using the Illumina HumanOmniExpressExome-8v1 arrays at the SciLifeLab NGI facility in Uppsala, Sweden. Fecal DNA was extracted from samples kept at -80°C using Qiagen QIAamp DNA Stool Mini Kits and analyzed using 16S rRNA gene amplicon sequencing (in the V1-V2 hypervariable region). This was performed on the Illumina MiSeq platform at the Institute of Clinical Molecular Biology (IKMB) in Kiel, Germany. After data merging and QC, we used data from 134 individuals (83 females, 51 males, mean age 54.8(11.3) yrs) in the microbiome GWAS. Of these, six PopCol participants were PPI users and 12 used antibiotics. The study was approved by the local Committee of Research Ethics (Forskningskommitté Syd) at Karolinska Institutet, Stockholm, in November 2001. Written informed consent was obtained from all participants.

RS (Rotterdam Study III)

The RS is a prospective population-based cohort study established in 1990 to study determinants of disease and disability in Dutch adult/elderly individuals aged ≥40 years. The original design and updates of this study have been described in detail[95]. The RS consists of four sub-cohorts and comprises approximately 18,000 inhabitants of the Ommoord, a suburb of Rotterdam, the Netherlands. In the current microbiome GWAS, data from the Rotterdam Study III have been used. Genotyping was performed using the Illumina HumanHap 550K and 610K. The collection of fecal samples started in 2012 and includes 3,932 participants. Fecal DNA was extracted using DiaSorin Arrow DNA (Isogen Life Science, De Meern, the Netherlands) with a bead-beating step. Sequencing of bacterial 16S gene, domain V3-V4, was performed in the Laboratory of Human Genetics at Erasmus MC Rotterdam, using the Illumina MiSeq platform[89]. After stringent QC, the overlap between samples with genotyping and microbial 16S sequencing data yielded 1,220 samples (705 females, 515 males, mean age 57 (5.9) yrs) for use in the microbiome GWAS analysis. Of these, 260 participants used proton pump inhibitors and none used antibiotics in the six months before fecal collection.

The study was approved by the IRB (Medical Ethics Committee) of the Erasmus Medical Center and by the review board of the Netherlands Ministry of Health, Welfare and Sports. Written informed consent was obtained from all participants.

SHIP (Study of Health in Pomerania)

The SHIP is a prospective longitudinal population-based cohort study encompassing two independent cohorts: SHIP (N=4,308; baseline examinations 1997-2001) and SHIP-TREND (N=4,420; baseline examinations 2008-2012)[96]. Individuals were invited to the SHIP study center for computer-assisted personal interviews and extensive examinations. Follow-up investigations are scheduled at 5-year intervals and have already been performed three times for SHIP and once for SHIP-TREND. For the microbiome GWAS project, data from the second SHIP wave (SHIP-2, 2008-2012) and the initial recruitment phase of SHIP-TREND were used. Genotyping was performed using the Genome-Wide Human SNP Array 6.0 (Affymetrix, Santa Clara, CA, USA) or the Infinium HumanOmni2.5 BeadChip (Illumina, San Diego, CA, USA) for SHIP and SHIP-TREND, respectively. Isolation of fecal DNA was done using the PSP Spin Stool DNA Kit (Stratec Biomedical AG, Birkenfeld, Germany). Fecal microbiota composition was determined based on the V1-V2 regions of the 16S rRNA gene on a MiSeq platform (Illumina) at the Institute of Clinical Molecular Biology (Christian Albrechts University of Kiel, Germany), as described before[97]. After comprehensive QC, 1,901 datasets (1,043 females, 858 males, mean age 53.7(14.0) yrs) with overlapping genotype and microbiome data were included in the current study. Of these, 149 individuals used PPIs and 25 had antibiotics at the time of inclusion.

The study was approved by the medical ethics committee of the University of Greifswald. Written informed consent was obtained from all participants.

HCHS/SOL (Hispanic Community Health Study/Study of Latinos)

The HCHS/SOL is a prospective, population-based cohort study of 16,415 Hispanic/Latino adults (ages 18-74 years) who were selected using a two-stage probability sampling design from four US communities (Chicago, IL; Miami, FL; Bronx, NY; San Diego, CA)[98,99]. The genotyping of this cohort was performed with an Illumina custom array (15041502 B3), which consists of the Illumina Omni 2.5M array (HumanOmni2.5-8v1-1) plus ~150k custom SNPs, with the QC performed at the HCHS/SOL Genetic Analysis Center[100]. Stool samples were collected in the HCHS/SOL Gut Origins of Latino Diabetes (GOLD) ancillary study, which enrolled participants from the HCHS/SOL approximately concurrently with the second visit for HCHS/SOL. Fecal DNA was extracted with the Qiagen MagAttract PowerSoil DNA kit with both chemical and physical (i.e. bead-beating) means to release DNA, as described in Marotz et al[101]. Sequencing of bacterial 16S gene, domain V4, was performed in Rob Knight's lab at the University of California San Diego (San Diego, CA, USA) using the Illumina MiSeq platform. After stringent QC, the overlap between genetically unrelated subjects with microbial 16S sequencing data yielded 1,097 samples (676 females, 421 males, mean age 57.2(10.9) yrs) used in the microbiome GWAS analysis. Of these, 341 used medication including PPI for indigestion, heartburn, or stomach problems and 321 used antibiotics in the six months before the fecal collection.

The study was approved by the Ethics and Institutional Review Boards of all institutions involved (Bronx Field Center – Albert Einstein School of Medicine; Chicago Field Center – University of Illinois Chicago; Miami Field Center – University of Miami; San Diego Field Center – San Diego State University). Written informed consent was obtained from all participants.

TwinsUK

TwinsUK is a population-based cohort established in 1992 to study the genetic and environmental basis of a range of complex diseases and conditions in adult/elderly twins from the UK[102]. Genotyping was performed using HumanHap610Q on 5,654 volunteers, followed by imputation. Fecal samples were collected between 2010 and 2016 for 1,793 of the genotyped twins. DNA was extracted using PowerSoil - htp DNA isolation kit and the V4 region of the 16s rRNA gene was sequenced using the Illumina MiSeq platform at the Department of Molecular Biology and Genetics, Cornell University, Ithaca, NY, USA. One twin out of each pair was randomly excluded from the population of 1,793 individuals, leaving 1,205 volunteers (1,101 females and 104 males, mean age 61.5(10.7) yrs) on which to conduct the microbiome GWAS analysis. Of these, 78 used PPIs and 62 had used antibiotics in the 6 months prior to sampling. The study was approved by the Cornell University IRB (Protocol ID 1108002388). Written informed consent was obtained from all participants.

#### Supplementary Note 2. Cohort Acknowledgements

CARDIA

The microbiome study was funded by K01-NH127159 (KAM), by the NIA Intramural Research Program (LJL), P30-DK056350 (KAM), and UL1-TR002489 (LJL). Genotyping was funded as part of the NHLBI Candidate-gene Association Resource (N01-HC-65226) and the NHGRI Gene Environment Association Studies (GENEVA) (U01-HG004729 (MF), U01-HG04424, and U01-HG004446). This manuscript has been reviewed and approved by CARDIA for scientific content.

COPSAC$_{2010}$

We express our deepest gratitude to the children and families of the COPSAC$_{2010}$ cohort study for all their support and commitment. We acknowledge and appreciate the unique efforts of the COPSAC research team, and Martin S. Mortensen at the University of Copenhagen for extracting and sequencing of microbial DNA. COPSAC is funded by private and public research funds, including from the Lundbeck Foundation (grant no. R16-A1694 to RB); the Danish Ministry of Health (grant no. 903516 to RB); the Danish Council for Strategic Research (grant no. 0603-00280B to RB); and the Capital Region Research Foundation. No pharmaceutical company was involved in the study.

DanFunD

The DanFunD staff are thanked for their commitment and perseverance in phenotyping study participants. We are grateful to Annemette Forman and Tine Lorentzen, lab technicians. The Lundbeck Foundation and the Tryg Foundation are thanked for their for financial support to the DanFunD microbiome study. The Novo Nordisk Foundation Center for Basic Metabolic Research is an independent Research Center at the University of Copenhagen, partially funded by an unrestricted donation from the Novo Nordisk Foundation (www.metabol.ku.dk). DB is receiving funding from NNF Copenhagen Bioscience PhD programme (NNF17CC0026760). TJ is receiving funding from TrygFonden [7-11-0213] and the Lundbeck Foundation [R155-2013-14070]. All other authors declare that there is no duality of interest associated with their contribution to this manuscript.

FGFP

We acknowledge the contribution of Flemish GPs and pharmacists to the data and sample collection. We thank all FGFP volunteers for participating in the project. The FGFP was funded with support from the Flemish government (IWT130359 to JR), the Research Fund–Flanders (FWO) Odysseus program (G.0924.09 to JR), the King Baudouin Foundation (2012-J80000-004 to JR), FP7 METACARDIS HEALTH-F4-2012-305312 to JR, VIB, the Rega Institute for Medical Research (Leuven), and KU Leuven. This study is partially funded by the FWO EOS program (30770923 to JR). MJ was funded by FWO through a Postdoctoral Fellowship (1299417N). JW is supported by the Strategic Priority Research Program of the Chinese Academy of Sciences (XDB29020000) and National Key Research and Development Program of China (2018YFC2000500). RB is funded by FWO through a Postdoctoral Fellowship (1221620N). DAH is supported by a Wellcome Trust Investigator Award (202802/Z/16/Z). KHW is supported by the Elizabeth Blackwell Institute for Health Research, University of Bristol, UK, and the Wellcome Trust Institutional Strategic Support Fund (204813/Z/16/Z). NJT is a Wellcome Trust Investigator (202802/Z/16/Z) supported by the University of Bristol NIHR Biomedical Research Centre (BRC-1215-20011) and works in the CRUK Integrative Cancer Epidemiology Program (C18281/A19169). NJT works in a unit that also receives funding from the MRC and the University of Bristol (MC_UU_12013/3, MC_UU_00011/6).

FOCUS/BSPSPC

We thank Ilona Urbach, Ines Wulf and Tonio Hauptmann of the IKMB microbiome laboratory and the staff of the IKMB sequencing facilities for excellent technical support. This work was supported by the Deutsche Forschungsgemeinschaft (DFG) Cluster of Excellence "Precision Medicine in Chronic Inflammation" (PMI, EXC2167, to AF), the Collaborative Research Center 1182, Origin and Function of Metaorganisms (ID: CRC1182, to AF).

GEM

All authors disclose no potential conflicts (financial, professional, or personal) that are relevant to the manuscript. KC is recipient of a Canada Research Chair in Inflammatory Bowel Diseases. KC receive funding to support this study from Crohn’s and Colitis Canada Grant #CCC-GEMIII, Canadian Institutes of Health Research (CIHR) Grant #CMF108031 and the Helmsley Charitable Trust. WT is a recipient of a Postdoctoral Fellowship Research Award from the CIHR Fellowship/ Canadian Association of Gastroenterology (CAG)/ Ferring Pharmaceuticals Inc. and a fellowship from the Department of Medicine, Mount Sinai Hospital, Toronto. JARG holds a

fellowship from the Department of Medicine, Mount Sinai Hospital, Toronto.
We thank the members of the CCC GEM Global Project Office: Ashleigh Goethel, Heather MacAulay, Stephanie Gosselin, Yurie Yamagishi, Clarissa Ratjen and all past team members for administrative support. The CCC GEM Project Research Consortium is composed of: Maria Abreu, Guy Aumais, Paul Beck, Charles Bernstein, Kenneth Croitoru, Leo Dieleman, Brian Feagan, Anne M Griffiths, David Guttman, Kevan Jacobson, Denis O. Krause*, Karen Madsen, John Marshall, Paul Moayyedi, Mark Ropeleski, Ernest Seidman*, Mark Silverberg, Scott Snapper, Andy Stadnyk, Hillary Steinhart, Michael Surette, Dan Turner, Thomas Walters, Bruce Vallance, Alain Bitton, Maria Cino, and Robert Baldassano, Lee Denson, Colette Deslandres, Wael El-Matary, Charlotte Hedin, Hans Herfarth, Peter Higgins, Seamus Hussey, Hien Huynh, Jeff Hyams, David Keljo, David Kevans, Charlie Lees, David Mack, Jerry McGrath, Sanjay Murthy, Anthony Otley, Remo Panaccione, Nimisha Parekh, Sophie Plamondon, Graham Radford-Smith, Joel Rosh, David Rubin, Michael Schultz, Corey Siegel, Martha Dirks, Herbert Brill, Sonia Michail, Hillel Noan, Jennifer Jones, Pierre Pare, Neil Leleiko, Fred Saibil, Dennis Cvitkovitch. (*deceased)

Generation R

We gratefully acknowledge the contribution of children and parents, general practitioners, hospitals, midwives and pharmacies in Rotterdam. The generation and management of GWAS genotype data for the Generation R Study was done at the Genetic Laboratory of the Department of Internal Medicine, Erasmus MC, the Netherlands. We thank Pascal Arp, Mila Jhamai, Marijn Verkerk, Lizbeth Herrera and Marjolein Peters for their help in creating, managing and QC of the GWAS database. The musculoskeletal research of the Generation R Study is partly supported by the European Commission grant HEALTH-F2-2008-201865-GEFOS to AGU and FR. The general design of Generation R Study is made possible by financial support from the Erasmus Medical Center, Rotterdam, the Erasmus University Rotterdam, the Netherlands Organization for Health Research and Development (ZonMw), the Netherlands Organization for Scientific Research (NWO), the Netherlands' Ministries of Health, Welfare and Sport and of Youth and Families. Additionally, the Netherlands Organization for Health Research and Development supported authors of this manuscript (ZonMw 907.00303, ZonMw 916.10159, ZonMw VIDI 016.136.361 to VWJ, and ZonMw VIDI 016.136.367to FR and supporting CMG). VWVJ received a Consolidator Grant from the European Research Council (ERC-2014-CoG-648916).

The study was also supported by funding from the EU's Horizon 2020 research and innovation program (733206, LIFECYCLE to VWJ). The generation and management of stool microbiome data for the Generation R Study was executed by the Human Genotyping Facility of the Genetic Laboratory in the Department of Internal Medicine, Erasmus MC, Rotterdam, the Netherlands. We thank Nahid El Faquir and Jolande Verkroost-Van Heemst for their help in sample collection and registration, and Kamal Arabe, Hedayat Razawy and Karan Singh Asra for their help in DNA isolation and sequencing. Furthermore, we thank Jeroen Raes and Jun Wang (KU Leuven, Belgium) for their guidance in 16S rRNA profiling and dataset generation. DJR was funded by an Erasmus MC mRACE grant "Profiling of the human gut microbiome".

KSCS

This study was supported by Basic Science Research Program through the National Research Foundation of Korea (NRF) funded by the Ministry of Education (NRF-2018R1D1A1B07050067 to HLK) and by the Medical Research Funds from Kangbuk Samsung Hospital. We thank the participants and staff of the Kangbuk Samsung Cohort Study for their collaboration, especially Yoosoo Chang and Seungho Ryu. The study was approved by the EUMC review board 2014-06-024 and KBSMC review board 2013-01-245. The computing resources were supported by the Global Science Experimental Data Hub Center (GSDC) Project and Korea Research Environment Open NETwork (KREONET) at the Korea Institute of Science and Technology Information (KISTI).

LLD

LifeLines DEEP: we thank participants and staff of the LifeLines DEEP cohort for their collaboration. We thank J. Dekens, M. Platteel, A. Maatman and J. Arends for management and technical support. This project was funded by the Netherlands Heart Foundation (IN-CONTROL CVON, grant no. 2012-03 to AZ and JF); by NWO (nos. NWO-VIDI 864.13.013 to JF, NWO-VIDI 016.178.056 to AZ, and NWO Gravitation Program ExposomeNL to AK and AZ), and by the European Research Council Starting Grant no. 715772 to AZ, who also holds Rosalind Franklin Fellowship from the University of Groningen. AD was supported by a VENI grant from NWO, H2020-SC1-HBC-28-2019-LONGITOOLS and WCRF-2017/1641. DVZ was supported by a VENI grant from NWO no. 194.006.

MIBS

We thank all participants and M. Hesselink for her support in data collection. Part of the data collection and analyses was funded by the Top Institute for Food and Nutrition (TIFN), Wageningen, the Netherlands.

METSIM

We thank all METSIM study participants. This study was funded by the Estonian Research Council grant (PUT1371 to EO), the EMBO Installation grant (No 3573 to EO), the US National Institutes of Health (NIH) grants (HL28481, HL144651 and DK117850 to AJL), the Academy of Finland (321428 to ML), Sigrid Juselius Foundation, Finnish Foundation for Cardiovascular Research, and the Centre of Excellence of Cardiovascular and Metabolic Diseases, supported by the Academy of Finland.

NGRC

We thank the NeuroGenetics Research Consortium (NGRC) participants and our collaborators S. Factor, C. Zabetian and E. Molho. Funding from the National Institutes of Health Grant 01NS036960 (to HP) (USA), Michael J Fox Foundation for Parkinson's Disease Research Edmond J. Safra Global Genetic Consortia (to HP), and NIH Training Grant T32NS095775 (to ZDW).

NTR

Funding was obtained from the Netherlands Organization for Scientific Research (NWO) and The Netherlands Organization for Health Research and Development (ZonMW) grants 904-61-090, 985-10-002, 912-10-020, 904-61-193,480-04-004, 463-06-001, 451-04-034, 400-05-717, Addiction-31160008, 016-115-035, 481-08-011, 400-07-080, 056-32-010, Middelgroot-911-09-032, OCW_NWO Gravity program –024.001.003, NWO-Groot 480-15-001/674, Center for Medical Systems Biology (CSMB, NWO Genomics), NBIC/BioAssist/RK(2008.024), Biobanking and Biomolecular Resources Research Infrastructure (BBMRI –NL, 184.021.007 and 184.033.111), X-Omics 184-034-019; Spinozapremie (NWO- 56-464-14192), KNAW Academy Professor Award (PAH/6635) and University Research Fellow grant (URF) to DIB; Amsterdam Public Health research institute (former EMGO+) , Neuroscience Amsterdam research institute (former NCA) ; the European Community's Fifth and Seventh Framework Program (FP5- LIFE QUALITY-CT-2002-2006, FP7- HEALTH-F4-2007-2013, grant 01254: GenomEUtwin, grant 01413: ENGAGE and grant 602768: ACTION); the European Research

Council (ERC Starting 284167, ERC Consolidator 771057, ERC Advanced 230374), Rutgers University Cell and DNA Repository (NIMH U24 MH068457-06), the National Institutes of Health (NIH, R01D0042157-01A1, R01MH58799-03, MH081802, DA018673, R01 DK092127-04, Grand Opportunity grants 1RC2 MH089951, and 1RC2 MH089995); the Avera Institute for Human Genetics, Sioux Falls, South Dakota (USA). Part of the genotyping and analyses were funded by the Genetic Association Information Network (GAIN) of the Foundation for the National Institutes of Health. Computing was supported by NWO through grant 2018/EW/00408559, BiG Grid, the Dutch e-Science Grid and SURFSARA.

PNP

We thank the Segal and Elinav group members for discussions. ES is supported by the Crown Human, Genome Center and the Else Kroener Fresenius Foundation; DL Schwarz; JN Halpern; L Steinberg; and grants from the European Research Council and the Israel Science Foundation. DR received a Levi Eshkol PhD Scholarship for Personalized Medicine from the Israeli Ministry of Science.

POPCOL

PopCol data generation and analysis has been partially supported by funding from the Swedish Research Council (VR project 2017-02403) available to MDA.

Rotterdam Study III

The generation and management of GWAS genotype data for the Rotterdam Study is supported by the Netherlands Organization of Scientific Research (NWO) Investments (no. 175.010.2005.011, 911-03-012 to AGU). This study was funded by the Research Institute for Diseases in the Elderly (014-93-015; RIDE2 to AGU), the Netherlands Genomics Initiative (NGI)/Netherlands Organization for Scientific Research (NWO) project no. 050-060-810 to AGU. We thank Pascal Arp, Mila Jhamai, Marijn Verkerk, Lizbeth Herrera, Marjolein Peters and Carolina Medina-Gomez for help in creating the GWAS database, and Linda Broer and Carolina Medina-Gomez for the creation and analysis of imputed data. The Rotterdam Study is funded by Erasmus Medical Center and Erasmus University, Rotterdam; the Netherlands Organization for Health Research and Development (ZonMw); the Research Institute for Diseases in the Elderly (RIDE); the Netherlands' Ministries of Education, Culture and Science, and of Health, Welfare and Sports; the European Commission (DG XII), and the Municipality of

Rotterdam. The authors thank the study participants, the Rotterdam Study staff, and the general practitioners and pharmacists who participated. The generation and management of stool microbiome data for the Rotterdam Study (RSIII-2) was executed by the Human Genotyping Facility of the Genetic Laboratory, Department of Internal Medicine, Erasmus MC, Rotterdam, the Netherlands. We thank Nahid El Faquir and Jolande Verkroost-Van Heemst for their help in sample collection and registration, and Kamal Arabe, Hedayat Razawy and Karan Singh Asra for help in DNA isolation and sequencing. Furthermore, we thank Jeroen Raes and Jun Wang (KU Leuven, Belgium) for guidance in 16S rRNA profiling and dataset generation. DJR was funded by an Erasmus MC mRACE grant "Profiling of the human gut microbiome".

SHIP

SHIP is part of the Community Medicine Research network of the University of Greifswald, Germany, which is funded by the Federal Ministry of Education and Research, the Ministry of Cultural Affairs, the Social Ministry of the Federal State of Mecklenburg-West Pomerania, and the network 'Greifswald Approach to Individualized Medicine (GANI_MED)', funded by the Federal Ministry of Education and Research (grant 03IS2061A to MML, HV, and UV). Genome-wide SNV typing data have been supported by the Federal Ministry of Education and Research (grant no. 03ZIK012 to GH) and a joint grant from Siemens Healthineers, Erlangen, Germany and the Federal State of Mecklenburg-West Pomerania. Further support was received from the RESPONSE-project (BMBF grant number 03ZZ0921E to MML), PePPP-project (ESF/14-BM-A55_0045/16 to MML), and EnErGie-project (ESF/14-BM-A55-0010/18 to MML).

HCHS/SOL

The Hispanic Community Health Study/Study of Latinos is a collaborative study supported by contracts from the US National Heart, Lung, and Blood Institute (NHLBI) to the University of North Carolina (HHSN268201300001I / N01-HC-65233 to RCK), University of Miami (HHSN268201300004I / N01-HC-65234 to RCK), Albert Einstein College of Medicine (New York), (HHSN268201300002I / N01-HC-65235 to RCK), University of Illinois at Chicago – HHSN268201300003I / N01-HC-65236 Northwestern Univ. to RCK), and San Diego State University (HHSN268201300005I / N01-HC-65237 to RCK). The following US organizations have contributed to the HCHS/SOL by transferring funds to the NHLBI: National Institute on Minority Health and Health Disparities, National Institute on Deafness and Other

Communication Disorders, National Institute of Dental and Craniofacial Research, National Institute of Diabetes and Digestive and Kidney Diseases, National Institute of Neurological Disorders and Stroke, NIH Institution-Office of Dietary Supplements. The Genetic Analysis Center at the University of Washington was supported by NHLBI and NIDCR contracts (HHSN268201300005C AM03 and MOD03 to RCK). Our study was also supported by grants from NIMHD (1R01MD011389-01 to RCK) and NHLBI (1R01HL140976-01 to RCK).

TwinsUK

TwinsUK is funded by the Welcome Trust, Medical Research Council, European Union, the National Institute for Health Research (NIHR)-funded BioResource, Clinical Research Facility and Biomedical Research Centre based at Guy's and St Thomas' NHS Foundation Trust in partnership with King's College London. SNP genotyping was performed by The Wellcome Trust Sanger Institute and National Eye Institute via NIH/CIDR. We thank Dr Julia K. Goodrich and Dr Ruth E. Ley and the Cornell technical team for generating the microbial data.

### Supplementary Note 3. Microbiome heterogeneity reduces the power of GWAS

The substantial variation in taxonomic composition driven by technical factors, including 16S domain and DNA extraction kits, has a significant effect on microbiome GWAS. For example, the genus *Bifidobacterium*, which showed the strongest genetic effect, was present in 93% of the samples in those cohorts that used the V4 domain of the 16S rRNA gene, but only 78% and 62% of the samples sequenced by V3-V4 and V1-V2 domains, respectively. Similar to the 16S domain, the DNA isolation method showed a strong influence on *Bifidobacterium* abundance, which ranged from 35.7% to 100% depending on the DNA isolation kit used (Table S3). Another example is the large effect of the sequencing domain on the presence of the Archaea, in particular genus *Methanobrevibacter*. The proportion of Archaea-positive individuals in cohorts sequenced by the V3-V4 or V4 domains was around 25–35%, similar to the prevalence estimated using shotgun metagenomics sequencing[2], whereas Archaea were not detected at all in cohorts that used the V1-V2 domain. This lack of Archaea detection dramatically reduces the sample size for mbTL mapping and may well explain the lack of genome-wide significant mbTLs for this domain, despite its moderate heritability ($H^2$=0.319). In general, half of the bacterial taxa that passed either the quantitative or binary mbTL filtering cutoff showed substantial differences in abundance or presence between the 16S domains or the DNA extraction methods (Table S3). However, our design did not always allow us to distinguish the causes of heterogeneity since the methodological discrepancy overlapped with biological variance between cohorts, including ethnicity, age, BMI and study design. For example, most of the cohorts that used the V1-V2 16S domain had German ancestry, whereas the group of cohorts that used the V3-V4 domain was very diverse and included several non-European and multi-ethnic cohorts (Table S1). Despite the expected effects of microbiome heterogeneity on the heterogeneity of mbTLs effects, we did not observe this correlation for either genome-wide significant or suggestive mbTLs (Supplementary Fig. 5a).

The effect sizes of the leading SNPs at the 31 genome-wide significant loci were consistent across cohorts, with the exception of one mbQTL presenting heterogeneity (Cochran's Q $P<0.05$): the *LCT* association with phylum Actinobacteria and a cluster of related taxa (class

Actinobacteria, order Bifidobacteriales, family *Bifidobacteriaceae* and genus *Bifidobacterium)*. Overall, the taxa with smaller effective sample size showed smaller numbers of genome-wide significant ($P<5x10^{-8}$) and suggestive ($P<1x10^{-5}$) associated loci (Supplementary Fig. 5b,c). Thus, the microbiome heterogeneity reduced the power of analysis but did not induce heterogeneity of mbTL effects.

### Supplementary Note 4. mbTL highlights

Of the loci with an association that did not achieve the stringent study-wide threshold, but did pass the nominal genome-wide significance threshold, the strongest mbQTL included 66 SNPs located in the *UHRF1BP1L* locus (12q23.1) that associated with the *Streptococcus* genus and *Streptococcaceae* family (rs11110281, $P=2.58x10^{-9}$). Eight genes located in this locus were identified by FUMA as positional candidates, including the closest gene, *UHRF1BP1L,* which is expressed in adipose tissue, liver and skeletal muscle. None of these genes could be prioritized as a prominent functional candidate based on published data and co-expression networks[103]. In the LLD cohort, the *Streptococcus* genus and *Streptococcaceae* family were positively correlated with stool levels of inflammatory markers chromogranin A ($R_{Sp}=0.22$, $P_{adj}=1.89x10^{-7}$) and calprotectin ($R_{Sp}=0.16$, $P_{adj}=1.4x10^{-3}$) and with the intake of proton pump inhibitors ($R_{Sp}=0.21$, $P_{adj}=9.42x10^{-7}$) (Table S10).

In mbBTL analysis, *Turicibacter*, which was the most heritable taxon determined by the twin analysis, was associated with rs555115 ($P=3.34x10^{-8}$), which is located in *IGSF21*, an immunoglobulin superfamily gene. *Turicibacter* is associated with decreased stool frequency and higher tea intake in the LLD cohort (Table S10) and is negatively associated with smoking in the FGFP (Table S11). The genus *Anaerostipes* was observed to be linked with rs17319026 ($P=4.67x10^{-8}$), located in carboxylesterase 5A (*CES5A*), which is involved in xenobiotic metabolism. Finally, the prevalence of the *Lachnospiraceae* family was associated with SNPs located in the olfactory receptor family 1 subfamily F member 1 (*OR1F1*). Although no associations have been reported between this SNP or the bacteria and food-related phenotypes, this gene is one of the olfactory receptors that regulates the perception of smell, which, in turn, might influence food preferences.

### Supplementary Figure 1. Alpha diversity Manhattan and QQ plots

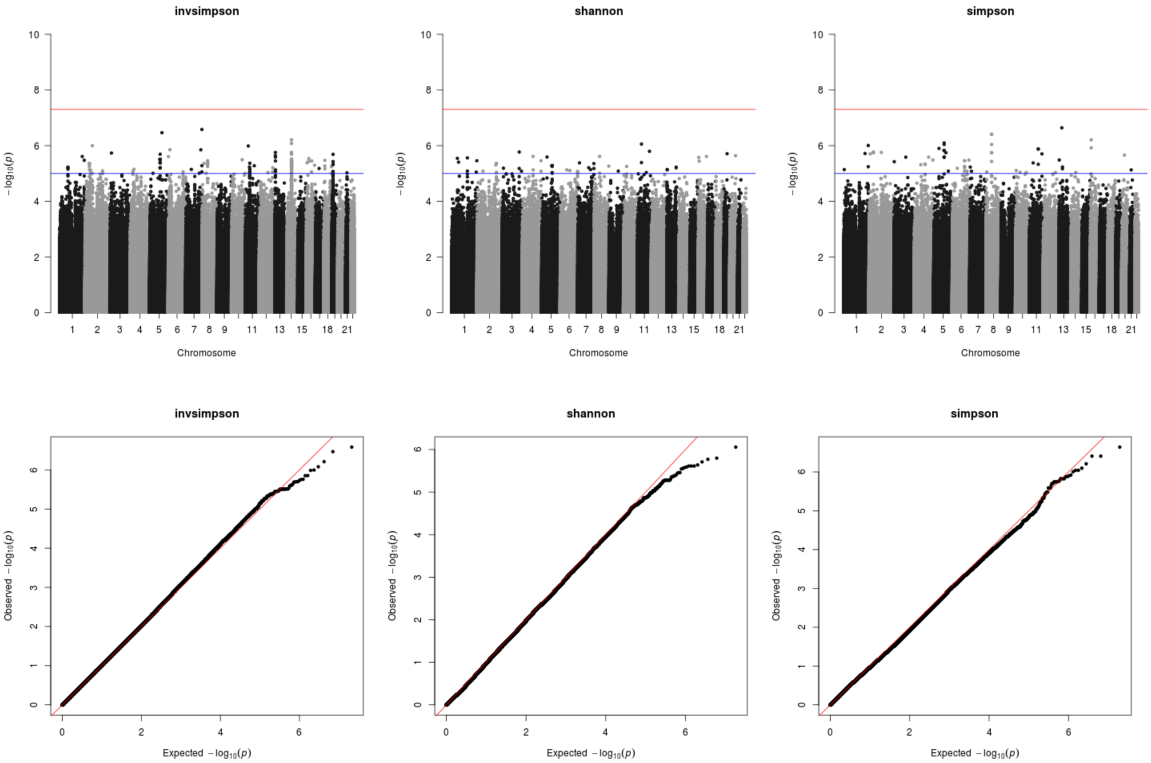


**Supplementary Figure 1**. Manhattan plots (top) and QQ plots (bottom) of GWAS on alpha diversity metrics (Inverse-Simpson, Shannon and Simpson indices). The name of the trait is given in the title of each plot. The Spearman correlation test (two-sided) was used to identify loci that affect the covariate-adjusted alpha diversity and SNP dosage. Blue horizontal line indicates suggestive genome-wide significance ($P=1x10^{-5}$).

### Supplementary Figure 2. Manhattan and QQ plots per mbQTL

**Supplementary Figure 2 (page 1).** Manhattan plots and QQ plots for mbQTLs (placed in an order of taxonomic level, from high to low). The name of the trait is given in the title of each plot. The Spearman correlation test (two-sided) was used to identify loci that affect the log-transformed, covariate-adjusted taxon abundance and SNP dosage. Samples with zero taxon abundance are excluded from the analysis. On the Manhattan plots, the red and blue horizontal lines indicate genome-wide and suggestive significance thresholds ($P=5x10^{-8}$ and $P=1x10^{-5}$, respectively).

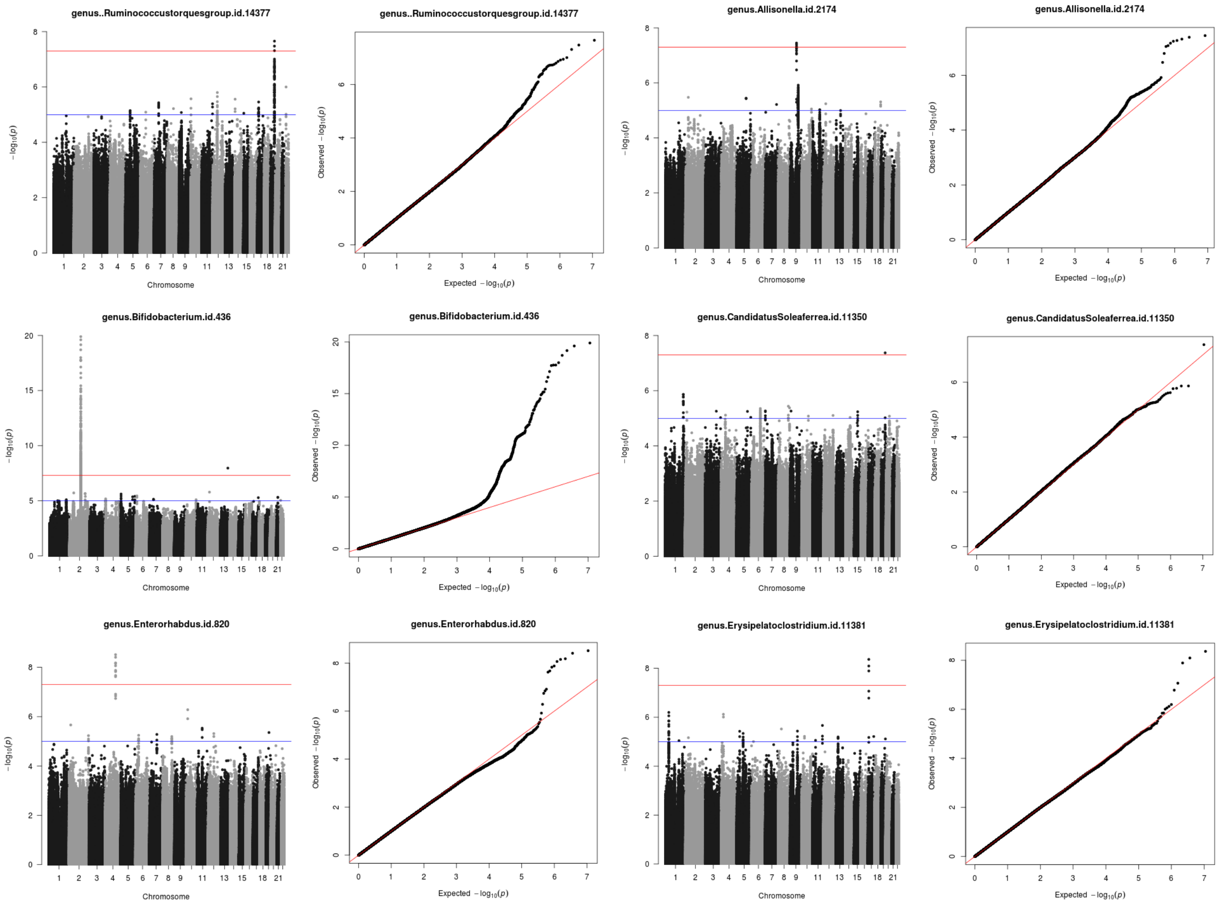

**Supplementary Figure 2 (page 2).** See legend on Supplementary Figure 2 page 1.

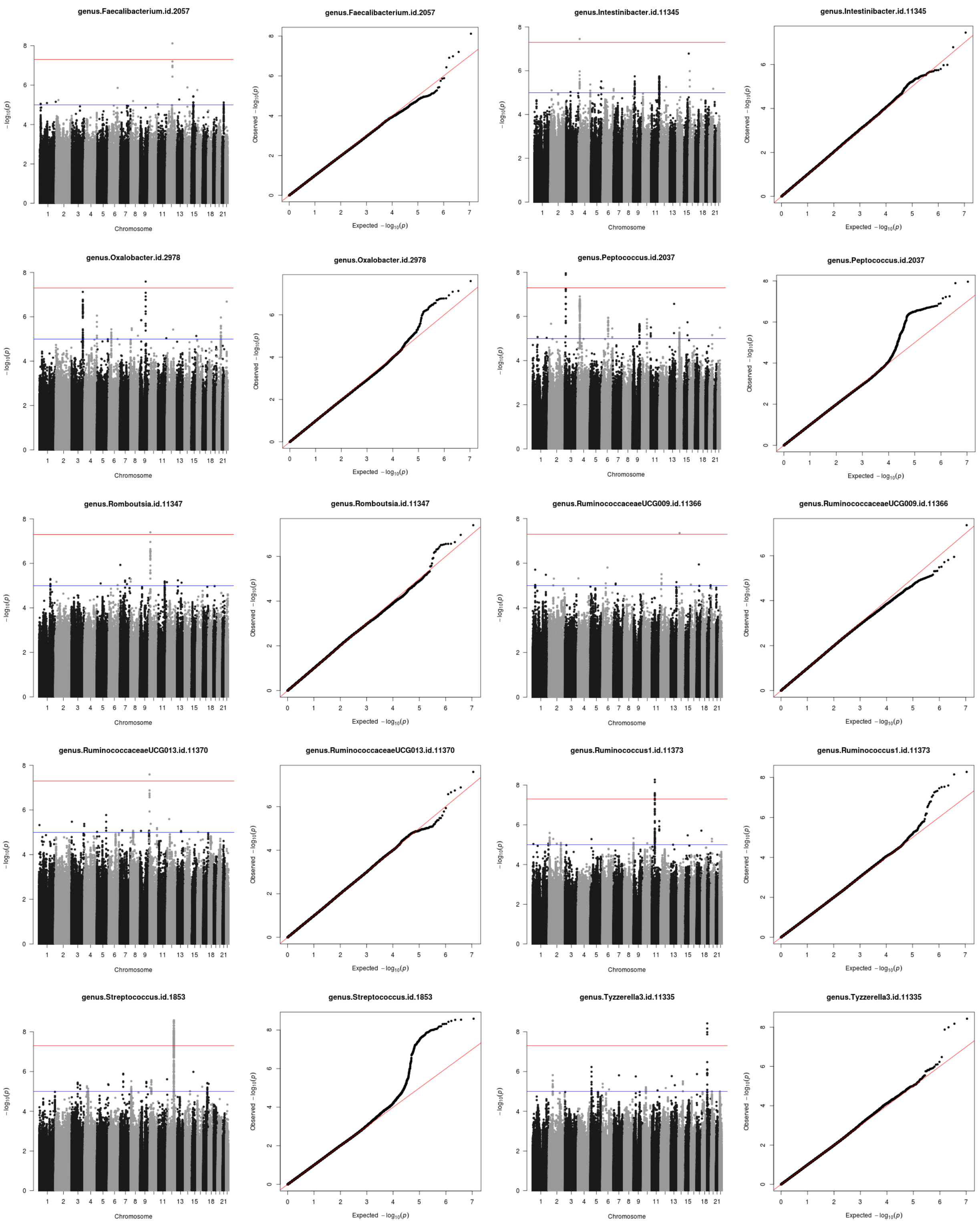

**Supplementary Figure 2 (page 3).** See legend on Supplementary Figure 2 page 1.

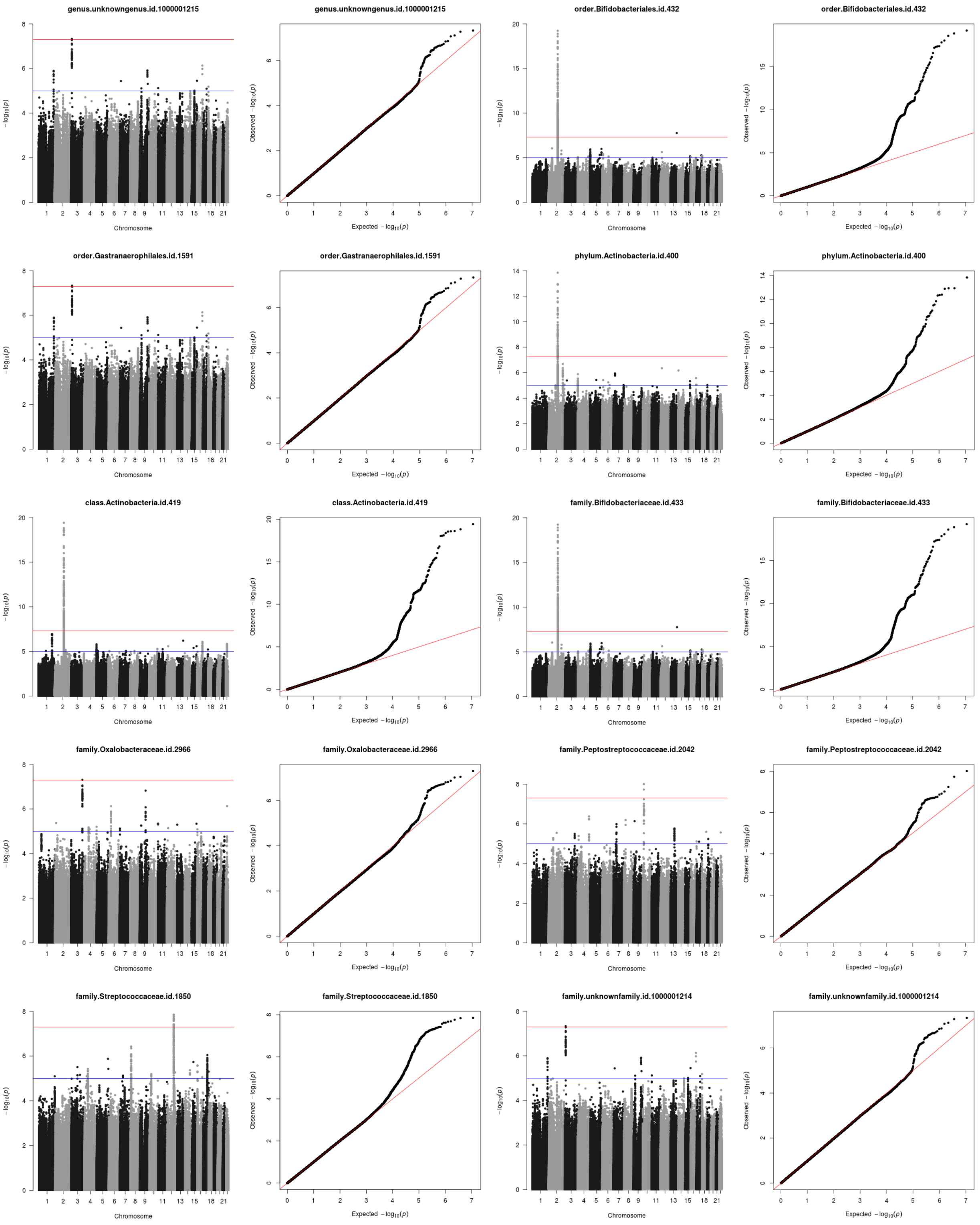

**Supplementary Figure 2 (page 4).** See legend on Supplementary Figure 2 page 1.

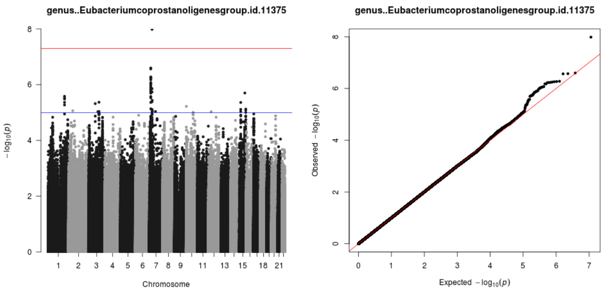

### Supplementary Figure 3. QQ plots for *Bifidobacterium* excluding *LCT* locus

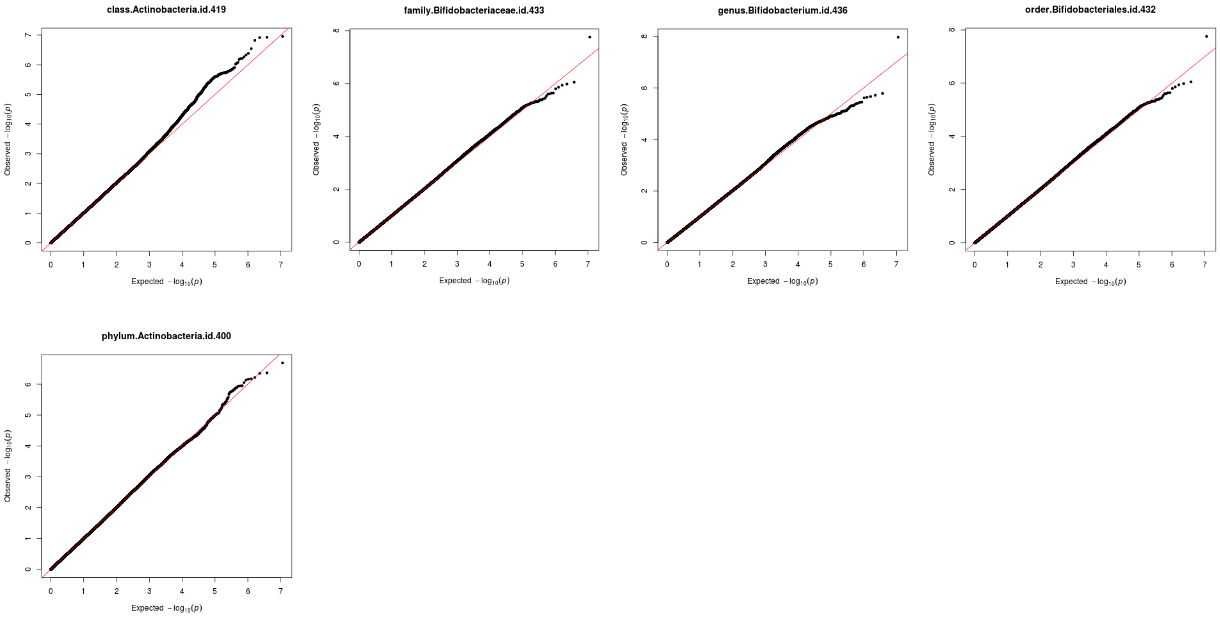


**Supplementary Figure 3.** QQplots of genus Bifidobacterium and its related upper level taxa, excluding the *LCT* locus (2MB upstream and downstream of the top SNP, rs182549). The Spearman correlation test (two-sided) was used to identify loci that affect the log-transformed, covariate-adjusted taxon abundance and SNP dosage. Samples with zero taxon abundance are excluded from the analysis.

### Supplementary Figure 4. Gene set enrichment analysis of mbQTLs

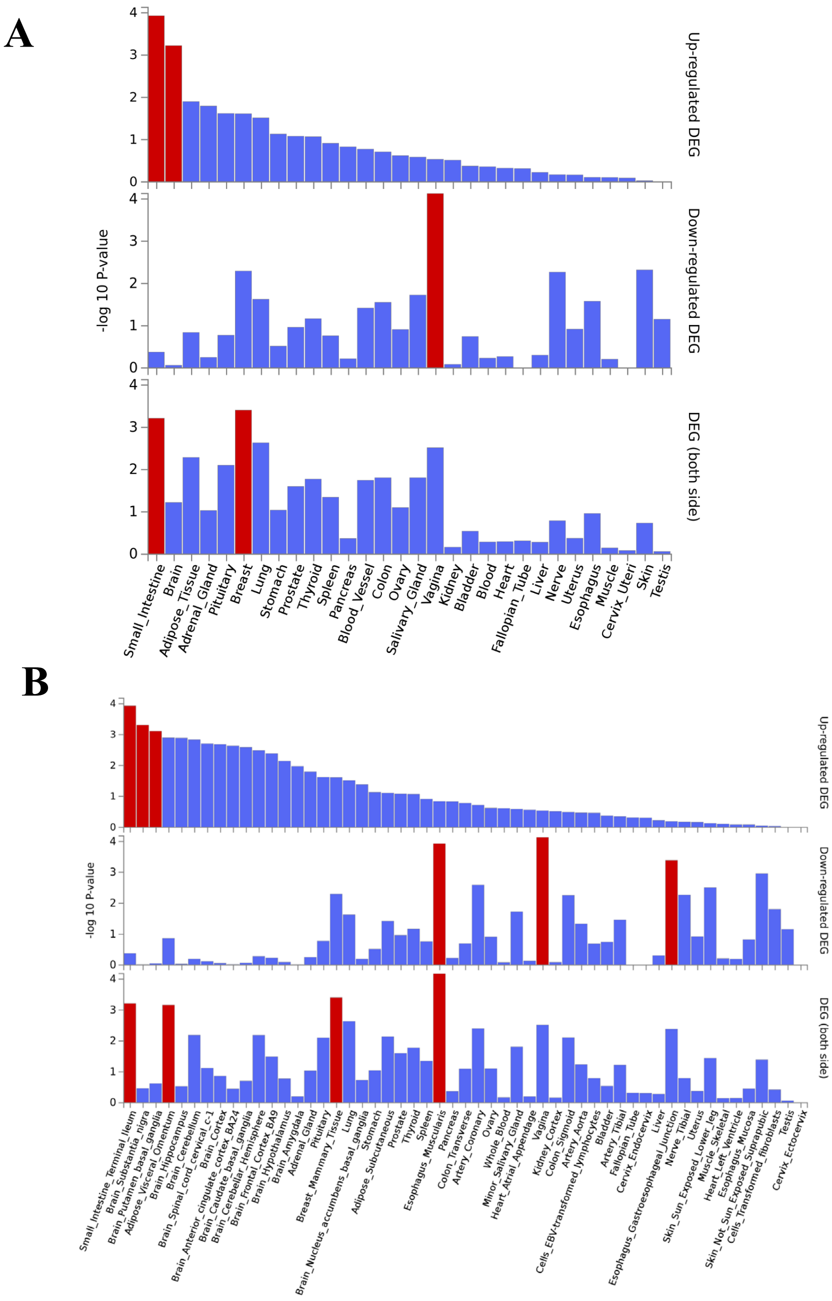


**Supplementary Figure 4.** Genome set enrichment analysis (GSEA) for mbQTLs. **(a)** GSEA analysis on 30 main tissue types. **(b)** GSEA analysis on 53 tissue types. DEG abbreviature means Differentially Expressed Genes

Supplementary Figure 5. Effect of taxon sample size and taxon inter-cohort variance on the detectability and heterogeneity of mbQTLs

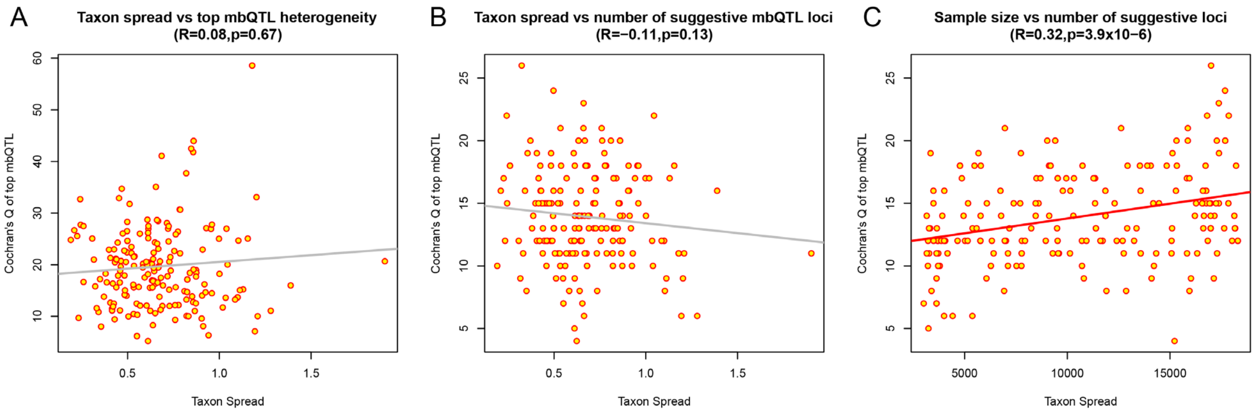


**Supplementary Figure 5.** Effect of taxon sample size and taxon inter-cohort variance on the detectability and heterogeneity of mbQTLs. **(a)** The correlation of taxon inter-cohort spread (calculated as $SD_{cohortMean}/E_{cohortMean}$) and Cochran's Q of top mbQTL per taxon. The genus *Bifidobacterium* and its related upper level taxa (family *Bifidobacterium*, phylum Bacteroidales, order Bacteroidia and class Actinobacteria) because there is a known biological origin for the heterogeneity. **(b)** The correlation of taxon inter-cohort spread (calculated as $SD_{cohortMean}/E_{cohortMean}$) with the number mbQTLs detected per taxon with a relaxed threshold of $P<10^{-5}$. The genus *Bifidobacterium* and its related upper-level taxa (family *Bifidobacterium*, phylum Bacteroidales, order Bacteroidia and class Actinobacteria) were excluded because there is a known biological origin for the heterogeneity. **(c)** The correlation of effective sample size (number of samples with non-zero bacterial abundance included in actual mbQTL analysis) with the number of detected mbQTL loci with at least one SNP with $P<10^{-5}$. Each point represents one taxon. A window of 1 mb was taken to define the locus**.**

### Supplementary Materials References
